# Pro-inflammatory macrophage activation does not require inhibition of mitochondrial respiration

**DOI:** 10.1101/2024.05.10.593451

**Authors:** Andréa B. Ball, Anthony E. Jones, Kaitlyn B. Nguyễn, Amy Rios, Nico Marx, Wei Yuan Hsieh, Krista Yang, Brandon R. Desousa, Kristen K.O. Kim, Michaela Veliova, Zena Marie del Mundo, Orian S. Shirihai, Cristiane Benincá, Linsey Stiles, Steven J. Bensinger, Ajit S. Divakaruni

**Affiliations:** Department of Molecular and Medical Pharmacology, University of California, Los Angeles, Los Angeles, CA, USA; Institute of Integrative Cell Biology and Physiology, Bioenergetics and Mitochondrial Dynamics Section, University of Münster, Schloßplatz 5, D-49078 Münster, Germany; Department of Microbiology, Immunology, and Molecular Genetics, University of California, Los Angeles, Los Angeles, CA, USA; Department of Medicine, University of California, Los Angeles, Los Angeles, CA, USA

## Abstract

Pro-inflammatory macrophage activation is a hallmark example of how mitochondria serve as signaling organelles. Upon classical macrophage activation, oxidative phosphorylation sharply decreases and mitochondria are repurposed to accumulate signals that amplify effector function. However, evidence is conflicting as to whether this collapse in respiration is essential or largely dispensable. Here we systematically examine this question and show that reduced oxidative phosphorylation is not required for pro-inflammatory macrophage activation. Only stimuli that engage both MyD88- and TRIF-linked pathways decrease mitochondrial respiration, and different pro-inflammatory stimuli have varying effects on other bioenergetic parameters. Additionally, pharmacologic and genetic models of electron transport chain inhibition show no direct link between respiration and pro-inflammatory activation. Studies in mouse and human macrophages also reveal accumulation of the signaling metabolites succinate and itaconate can occur independently of characteristic breaks in the TCA cycle. Finally, *in vivo* activation of peritoneal macrophages further demonstrates that a pro-inflammatory response can be elicited without reductions to oxidative phosphorylation. Taken together, the results suggest the conventional model of mitochondrial reprogramming upon macrophage activation is incomplete.

## INTRODUCTION

Metabolic alterations are tightly linked to macrophage function and fate^1–4^. Classical, pro-inflammatory activation triggered by exposure to lipopolysaccharide (LPS) causes a dramatic shift in macrophage energy metabolism: ATP production is shifted almost entirely to glycolysis, the TCA cycle is dramatically rewired, and oxidative phosphorylation is almost entirely inhibited^2,5,6^.

It is generally accepted that this respiratory inhibition and a shift to ‘aerobic glycolysis’ – largely due to excessive nitric oxide production^7–9^ – is an essential feature of pro-inflammatory activation^10–12^. Mitochondria are thought to be repurposed away from oxidative phosphorylation to generate metabolites and other mitochondrial signals that enhance macrophage function^13–16^. Genetic loss-of-function studies also point to a specific role for oxidative phosphorylation and respiratory chain function: myeloid-specific loss of a subunit of respiratory complex I reportedly enhances the pro-inflammatory phenotype whereas no phenotypic changes were observed upon myeloid-specific ablation of the mitochondrial pyruvate carrier^17,18^.

Other reports, however, have demonstrated preserved or even enhanced pro-inflammatory macrophage function under conditions where oxidative phosphorylation remains functional^7,9,19,20^. Additionally, as has been previously suggested, physiologically relevant mitochondrial signals such as redox changes, metabolite accumulation, or superoxide production from reverse electron transport (RET) do not require mitochondrial damage or dysfunction^21,22^. It therefore remains unclear to what extent this hallmark collapse in mitochondrial ATP production is a requisite, causal driver of macrophage effector function or simply an associated epiphenomenon.

Here we use pharmacologic, genetic, human, and *in vivo* models to systematically detail that this well-established reduction in oxidative phosphorylation is unexpectedly dispensable for the induction of the macrophage pro-inflammatory response. We show that (i) not all pro-inflammatory stimuli decrease mitochondrial respiration, (ii) the signaling metabolites itaconate and succinate can accumulate in mouse and human macrophages without characteristic ‘breaks’ in the TCA cycle, (iii) pharmacologic and genetic inhibition of the respiratory chain does not amplify the pro-inflammatory response, (iv) respiratory inhibition does not temporally align with the induction of the pro-inflammatory response, and (v) mouse peritoneal macrophages activated via sterile inflammation retain normal oxidative phosphorylation.

## RESULTS

### Not all pro-inflammatory stimuli elicit uniform bioenergetic responses

A frequently studied means to classically activate pro-inflammatory macrophages is use of lipopolysaccharide (LPS), an outer membrane component of Gram-negative bacteria. LPS is a Toll-like receptor 4 (TLR4) agonist. When offered at high concentrations, it can activate both the myeloid differentiation primary response protein 88 (MyD88) and TIR-domain-containing adaptor-inducing interferon-β (TRIF) adaptor proteins (Fig. 1A & S1A)^23^. Treatment of mouse bone marrow-derived macrophages (BMDMs) with 50 ng/mL LPS for 24 hr. resulted in decreased mitochondrial respiration, as is well established (Figs. 1B&C)^7,8,14,24^. Since TLR4 is upstream of MyD88 and TRIF, we sought to determine which signaling arm is responsible for the decrease in respiration. We therefore exposed BMDMs to Pam3CSK4 (Pam3; a TLR2 agonist which specifically activates MyD88) as well as polyinosinic-polycytidylic acid (Poly I:C; a TLR3 agonist which specifically activates TRIF) (Fig. 1D)^23,25^. Treatment with either Pam3 or Poly I:C for 24 hr. was sufficient to elicit the expression of characteristic pro-inflammatory genes and the secretion of inflammatory cytokines (Figs. S1B-D). However, neither Pam3 nor Poly I:C exposure caused the profound respiratory inhibition observed with LPS (Figs. 1E&F).

**Figure 1:**
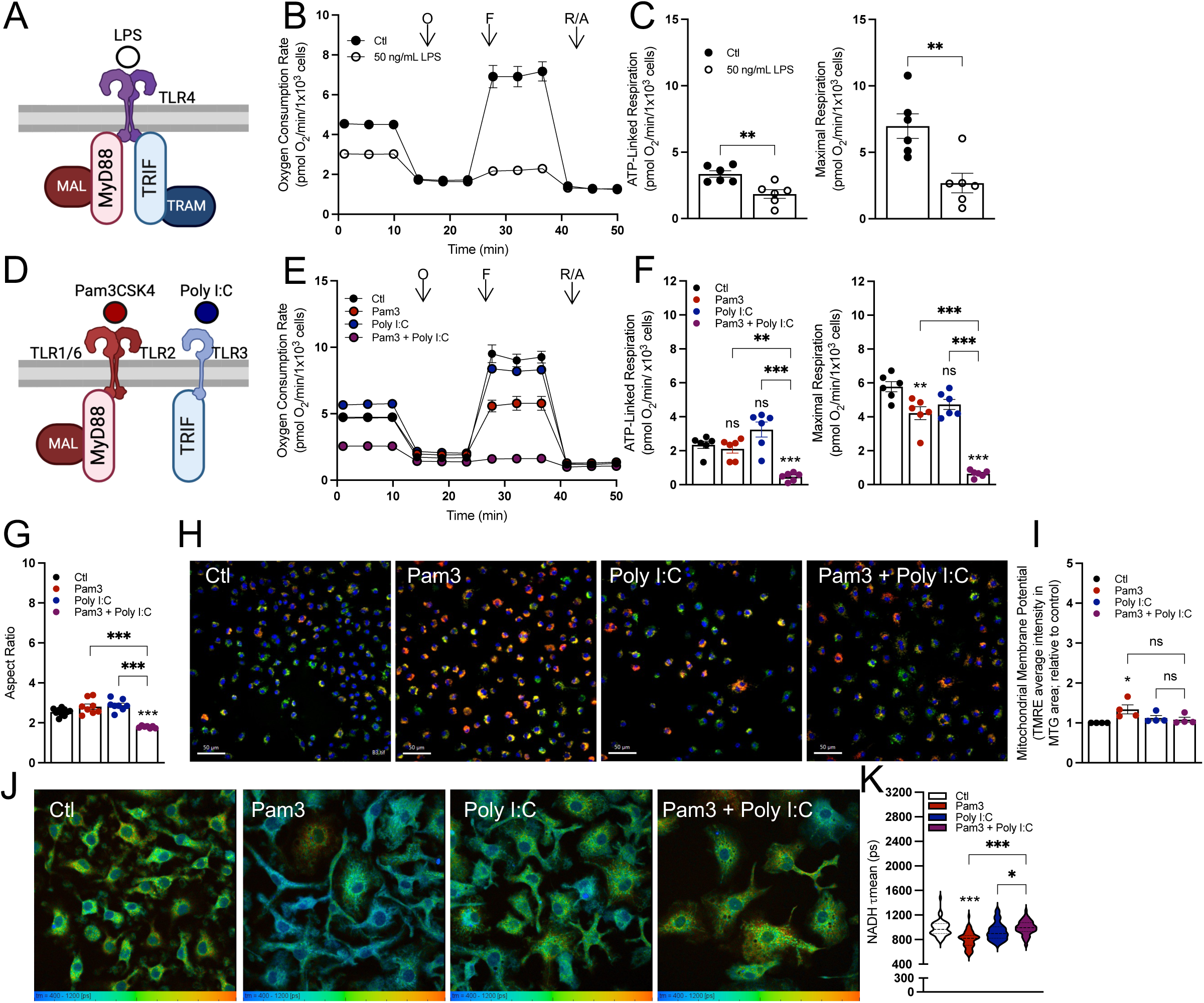
Differential effects on mitochondrial function from various pro-inflammatory stimuli after 24 hr. **A**) A graphical schematic of the initial adapter proteins downstream of TLR4 engagement by lipopolysaccharide (LPS). TLR4, Toll-like receptor 4; MyD88, myeloid differentiation primary response protein; MAL (also known as TIRAP), MyD88-adapter like; TRIF, TIR-domain-containing adapter-inducing interferon-β; TRAM, TRIF-related adapter molecule. **B**) Representative oxygen consumption trace with control BMDMs (Ctl) and BMDMs treated with 50 ng/mL LPS for 24 hr. Where not visible, error bars are obscured by the symbol. O, oligomycin; F, carbonyl cyanide-p-trifluoromethoxyphenylhydrazone (FCCP); R/A, rotenone/antimycin A (n = 1 biological replicate with 5 technical replicates). **C**) ATP-Linked and maximal respiration rates for control (Ctl) and BMDMs treated with 50 ng/mL LPS for 24 hr. (n = 6). **D**) A graphical schematic of the initial adapter proteins downstream of either TLR2 or TLR3 engagement by pathogen associated molecular patterns. TLR2, Toll-like receptor 2; TLR3, Toll-like receptor 3. **E**) The oxygen consumption rates from a representative experiment with control BMDMs (Ctl) and BMDMs activated with Pam3, Poly I:C, or Pam3 + Poly I:C for 24 hr. Where not visible, error bars are obscured by the symbol. O, oligomycin; F, carbonyl cyanide-p-trifluoromethoxyphenylhydrazone (FCCP); R/A, rotenone/antimycin A (n = 1 biological with 5 technical replicates). **F**) ATP-Linked and maximal respiration rates for control (Ctl) and BMDMs treated with Pam3, Poly I:C, or Pam3 + Poly I:C for 24 hr. (n = 6). **G**) Mitochondrial aspect ratio quantified using FIJI image analysis software for control (Ctl) and BMDMs treated with Pam3, Poly I:C, or Pam3 + Poly I:C for 24 hr. (images taken from n = 7-10 cells from three independent cell preparations). **H**) Representative images of control (Ctl) and BMDMs treated with Pam3, Poly I:C, or Pam3 + Poly I:C for 24 hr. Nuclei are stained with Hoechst, mitochondria are stained with MTG and membrane potential as determined by TMRE. **I**) Bulk membrane potential as measured by Tetramethylrhodamine, ethyl ester (TMRE) fluorescence per mitochondrial area detected by MitoTracker Green (MTG) for control (Ctl) and BMDMs treated with Pam3, Poly I:C, or Pam3 + Poly I:C for 24 hr. Data is shown relative to control (n = 4). **J**) Representative images of control (Ctl) and BMDMs treated with Pam3, Poly I:C, or Pam3 + Poly I:C for 24 hr. analyzed via fluorescence lifetime imaging (FLIM). **K**) Mean endogenous NADH lifetime (**τ**mean) for whole cell measured in picoseconds (ps) for control (Ctl) and BMDMs treated with Pam3, Poly I:C, or Pam3 + Poly I:C for 24 hr. Data is from one biological replicate with each data point representing one cell (n = 50-80). All data are mean ± SEM with statistical analysis conducted on data from biological replicates, each of which included multiple technical replicates, unless otherwise indicated. Statistical analysis for (**C**) was performed as an unpaired, two-tailed, t-test. Statistical analysis for (**F-I**) was performed as an ordinary one-way, ANOVA followed by Tukey’s *post hoc* multiple comparisons test.

We therefore hypothesized that a stimulus that engaged both MyD88 and TRIF was required to elicit this phenotype. Indeed, co-treatment with both Pam3 and Poly I:C was sufficient to substantially decrease mitochondrial respiration after 24 hr. (Figs. 1E&F). This relationship was also reproduced with Poly I:C and imiquimod (IMQ), a TLR7 agonist that engages MyD88 (Figs. S1E&F). The MyD88-linked agonists Pam3 and IMQ slightly decreased maximal respiration, though this was not nearly to same extent as when used in combination with Poly I:C (Figs. 1F, S1E). Furthermore, treatment with heat-killed *Staphylococcus aureus* (HKSA) – a physiologically relevant TLR2 agonist – yielded a similar phenotype. No decrease was observed in ATP-linked respiration and only a minimal defect was observed in maximal respiration (Fig. S1G).

We subsequently used other approaches including measurements of mitochondrial morphology, mitochondrial membrane potential, and NADH fluorescence lifetime imaging^26^ to determine whether this cooperativity between Pam3 and Poly I:C extended to other aspects of mitochondrial function. Similarly to respiration, neither ligand alone appreciably altered mitochondrial morphology after 24 hr. but co-treatment decreased the aspect ratio, suggesting increased fragmentation (Figs. 1G, S1H). Unexpectedly, we did not observe the same cooperativity with measurements of either mitochondrial membrane potential or NADH fluorescence lifetime imaging. 24 hr. treatment with Pam3 resulted in increased membrane potential relative to vehicle controls, whereas Poly I:C and the combination of both ligands did not cause a significant change (Figs. 1H&I). Mitochondrial membrane potential was measured as the average tetramethylrhodamine, ethyl ester (TMRE) intensity in area positive for MitoTracker Green (MTG) staining. Additionally, fluorescence lifetime imaging (FLIM)^27,28^ of total cellular NADH revealed a heterogeneous response in macrophages treated with Pam3, Poly I:C, or both ligands (Figs. 1J&K), with the most pronounced difference being a shorter lifetime in Pam3-treated macrophages relative to vehicle controls. The FLIM indicates the redox status and/or size of the macrophage pyridine nucleotide pool is subject to signal-specific remodeling during pro-inflammatory activation^27,28^. Collectively, the results in Fig. 1 show the bioenergetic response during classical macrophage activation is not uniform, and engagement of both the MyD88 and TRIF adaptor proteins is required to alter mitochondrial respiration and morphology.

Having used pharmacology to determine that engagement of both MyD88 and TRIF is required to decrease mitochondrial respiration, we sought to validate this with genetic proof-of-concept using BMDMs isolated from mice lacking either MyD88 or TRIF. If both adaptor proteins are required, then loss of either protein would rescue respiration in response to relevant stimuli. Indeed, the loss of either protein was sufficient to increase ATP-linked and maximal respiration in macrophages polarized for 24 hr. with stimuli that engage both MyD88 and TRIF: Pam3 with Poly I:C (Figs. 2A&B, S2A), 50 ng/mL LPS (Fig. 2C), and IMQ with Poly I:C (Figs. S2B&C).

**Figure 2:**
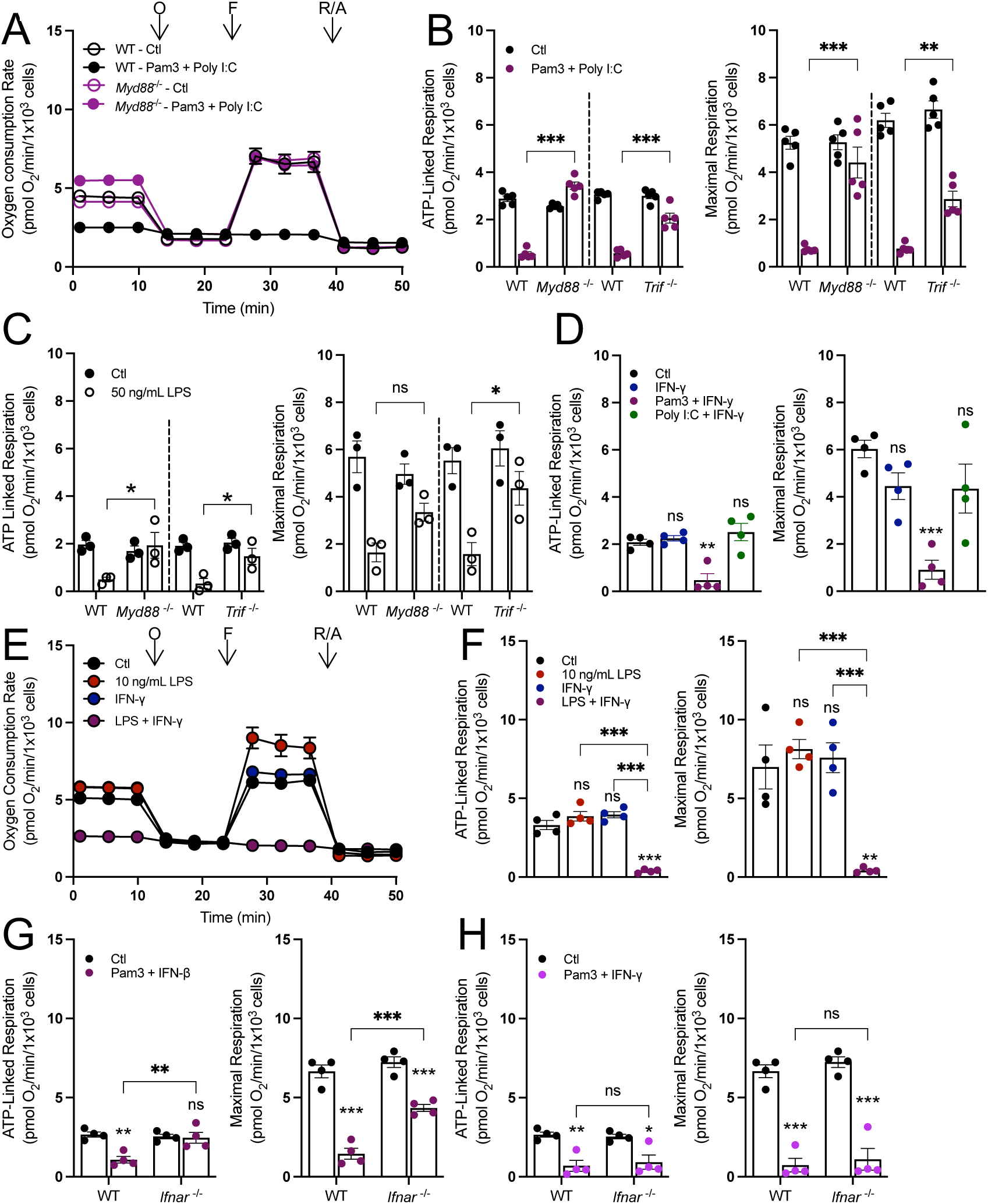
Both MyD88- and IFN-linked signaling are required to decreasing oxidative phosphorylation. **A**) Representative oxygen consumption trace with wildtype (WT) or MyD88-null (*Myd88*^-/-^) control (Ctl) and BMDMs treated with Pam3 + Poly I:C for 24 hr. Where not visible, error bars are obscured by the symbol. O, oligomycin; F, carbonyl cyanide-p-trifluoromethoxyphenylhydrazone (FCCP); R/A, rotenone/antimycin A (n = 1 biological replicate with 5 technical replicates). **B&C**) ATP-linked and maximal respiration rates for wildtype (WT), MyD88-null (*Myd88*^-/-^), and TRIF-null (*Trif*^-/-^) control (Ctl) and BMDMs activated with Pam3 + Poly I:C (n = 5) (**B**) or 50 ng/mL LPS for 24 hr. (n = 3) (**C**). **D**) ATP-linked and maximal respiration rates for control (Ctl) and BMDMs treated with IFN-γ, Pam3 + IFN-γ, or Poly I:C + IFN-γ (n = 4). **E&F**) Representative oxygen consumption rates, where not visible, error bars are obscured by the symbol. O, oligomycin; F, FCCP; R/A, rotenone/antimycin A (n = 1 biological with 5 technical replicates) (**E**) and ATP-linked and maximal respiration for control BMDMs (Ctl) and BMDMs activated with 10 ng/mL LPS, 20 ng/mL IFNgamma, or LPS + IFNgamma for 24 hr. (n=4) (**F**). **G&H**) ATP-linked and maximal respiration of WT and IFNAR-null (*Ifnar*^-/-^) control (Ctl) and BMDMs activated with Pam3 + IFNβ (n = 4) (**G**) or Pam3 + IFN-γ (n = 4) (**H**). All data are mean ± SEM with statistical analysis conducted on data from biological replicates, each of which included multiple technical replicates, unless otherwise indicated. Statistical analysis for (**B**), (**C**), (**G**), and (**H**) was performed as an ordinary two-way, ANOVA followed by Sídák’s *post hoc* multiple comparisons test. Statistical analysis for (**D**) and (**F**) was performed as an ordinary one-way, ANOVA followed by Tukey’s *post hoc* multiple comparisons test.

We then sought to better understand which signaling program downstream of TRIF is required to lower oxidative phosphorylation, hypothesizing that this was an interferon-linked response^23^. Indeed, treatment of BMDMs with IFN-γ in combination with Pam3, 10 ng/mL LPS, or HKSA for 24 hours collapsed mitochondrial respiration (Figs. 2D-F; S2D). No significant change in respiration was observed when treating macrophages with both Poly I:C and IFN-γ (Fig. 2D), and the effect of IFN-γ and 10 ng/mL LPS was lost in *Myd88^-/-^* BMDMs (Fig. S2E), further highlighting the requirement of MyD88. Additionally, macrophages isolated from mice lacking the Type I interferon receptor (IFNAR) had restored respiration upon co-treatment with Pam3 and IFN-β but not Pam3 and IFN-γ, a Type II interferon that bypasses IFNAR (Figs. 2G&H). Altogether, genetic data further show that reduced mitochondrial respiration is not a characteristic feature of all pro-inflammatory stimuli, but rather only those that engage both MyD88- and IFN-linked signaling.

Increased glycolysis is another bioenergetic hallmark of pro-inflammatory macrophage activation^6,29–32^. To better understand which signaling pathways were required to increase glycolysis, we treated BMDMs with Pam3, 10 ng/mL LPS, Poly I:C, or IFN-γ for 24 hr. Only stimuli upstream of MyD88 (Pam3 and 10 ng/mL LPS) substantially increased intracellular lactate abundance and rates of lactate efflux by 3-4-fold (Fig. S3A&B). This effect was lost in BMDMs lacking MyD88, establishing the requirement of MyD88-linked signaling for the profound increase in glycolysis (Fig. S3C). The results further reinforce the lack of a universal bioenergetic phenotype that characterizes all pro-inflammatory stimuli.

### Accumulation of pro-inflammatory metabolites is not linked to changes in mitochondrial respiration

Having established that not all pro-inflammatory stimuli collapse oxidative phosphorylation, we next aimed to define precisely which features of the pro-inflammatory response correlated with reduced oxygen consumption. We first applied this approach to study the accumulation of the TCA cycle-linked metabolites succinate and itaconate (Fig. 3A), hypothesizing that engagement of both MyD88 and TRIF would result in their enhanced accumulation relative to activating either pathway in isolation. During LPS ± IFN-γ activation, distinct ‘breaks’ occur in the TCA cycle, slowing activity of isocitrate dehydrogenase (IDH) and succinate dehydrogenase (SDH)^5,15,33^. It is generally accepted that this enzyme inhibition slows oxidative phosphorylation and facilitates the accumulation of metabolites itaconate and succinate, both of which can impact cell signaling, post-translational modifications, and the antimicrobial response^6,16,34,35^.

**Figure 3:**
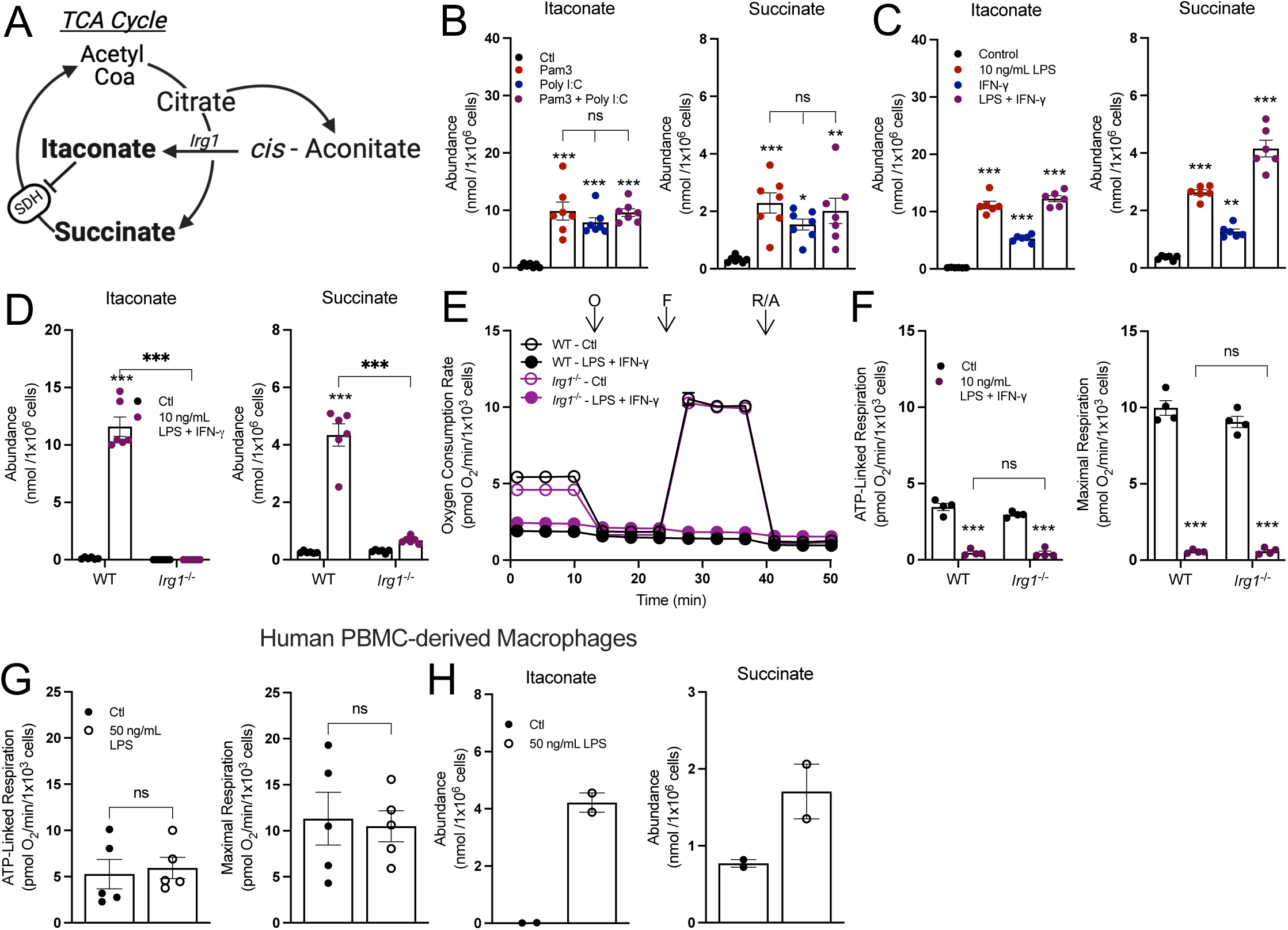
Accumulation of succinate and itaconate is independent of alterations to oxidative phosphorylation. **A**) A schematic depicting itaconate and succinate in the context of the tricarboxylic acid cycle (TCA cycle). IRG1, cis-aconitase decarboxylase 1; SDH, succinate dehydrogenase. **B**) Intracellular abundances of itaconate and succinate from control (Ctl) and BMDMs treated with Pam3, Poly I:C, or Pam3 + Poly I:C for 24 hr. (n = 7). **C**) Intracellular abundances of itaconate and succinate from control (Ctl) and BMDMs treated with 10 ng/mL LPS, 20 ng/mL IFN-γ, or LPS + IFN-γ for 24 hr. (n = 6). **D**) Intracellular abundances of itaconate and succinate in wildtype (WT) or IRG1-null (*Irg1*^-/-^) control (Ctl) and BMDMs treated with LPS + IFN-γ for 24 hr. (n = 6). **E&F**) Representative oxygen consumption rates, where not visible, error bars are obscured by the symbol, O, oligomycin; F, FCCP; R/A, rotenone/antimycin A (n = 1 biological with 5 technical replicates) and (**E**) ATP-linked and maximal respiration for WT or Irg1^-/-^ control (Ctl) and BMDMs treated with LPS + IFN-γ for 24 hr. (n = 4) (**F**). **G**) ATP-linked and maximal respiration rates for control (Ctl) and HMDMs treated with 50 ng/mL LPS for 24 hr. (n = 5). **H**) Intracellular abundances of itaconate and succinate from control (Ctl) and HMDMs treated with 50 ng/mL LPS for 24 hr. (n = 2). All data are mean ± SEM with statistical analysis conducted on data from biological replicates, each of which included multiple technical replicates, unless otherwise indicated. Statistical analysis for (**B**) and (**C**) was performed as an ordinary one-way, ANOVA followed by Tukey’s *post hoc* multiple comparisons test. Statistical analysis for (**D**) and (**F**) was performed as an ordinary two-way, ANOVA followed by Sídák’s *post hoc* multiple comparisons test. Statistical analysis for (**G**) was performed as an unpaired, two-tailed t-test.

Although neither Pam3 nor Poly I:C inhibit oxidative phosphorylation, steady-state levels of intracellular itaconate and succinate still accumulated substantially after 24 hr. treatment and at levels comparable with co-treatment (Fig. 3B). This pattern was reproduced with other pairs of MyD88- and IFN-linked ligands, such as 10 ng/mL LPS with IFN-γ (Fig. 3C) or IMQ with Poly I:C or (Fig. S4A).

Next, we studied BMDMs harvested from *Irg1^-/-^* mice to further understand the extent to which accumulation of succinate and itaconate is linked with the collapse in respiration from LPS ± IFN-γ. Siphoning carbon out of the TCA cycle to generate itaconate and the subsequent inhibition of SDH is thought to contribute to the restricted oxidative phosphorylation observed upon classical macrophage activation^14,15^. However, the loss of *Irg1* and the inability to accumulate itaconate and succinate had no effect on the respiratory inhibition observed in response to 24 hr. treatment with either LPS and IFN-γ (Figs. 3D-F) or Pam3 and Poly I:C. (Figs. S4B&C). Additionally, as has been previously reported^20^, human monocyte-derived macrophages (HMDMs) do not exhibit respiratory inhibition in response to stimulation with LPS (Fig. 3G). However, HMDMs readily accumulate itaconate and succinate (Fig. 3H). Altogether, the data show that macrophages do not need to repurpose mitochondria away from oxidative phosphorylation to directly support the generation of pro-inflammatory metabolites.

Since pro-inflammatory macrophages can accumulate signaling metabolites while maintaining oxidative phosphorylation, we sought to better understand the pathways by which this occurs. After 24 hr., citrate abundance sharply increased only upon co-treatment with Pam3 and Poly I:C and not with either ligand alone (Fig. 4A), confirming a ‘break’ in the TCA cycle and NO-mediated inhibition of IDH and aconitase activity only with joint activation of MyD88 and TRIF signaling^7^. To better examine individual enzymes and pathways, we measured oxygen consumption in plasma membrane-permeabilized BMDMs to directly offer mitochondria specific respiratory substrates (Fig. S5A). BMDMs were treated with Pam3, Poly I:C, or both ligands, and offered various substrate pairs in order to better understand enzymatic capacity in response to these stimuli. We observed the same pattern regardless of whether permeabilized cells were offered pyruvate with malate, glutamate with malate, succinate with rotenone, or citrate: only co-treatment with Pam3 and Poly I:C reduced oxygen consumption rates (Fig. 4B). The results, particularly with citrate-driven respiration, further indicate both a MyD88 and IFN signal are required for respiratory inhibition and reduced IDH activity. As treatment with Pam3 or Poly I:C alone can accumulate the SDH inhibitor itaconate (Fig. 3B), it was unexpected that succinate-driven respiration was unchanged. However, a time-course revealed this is likely due to exogenously added succinate outcompeting endogenous itaconate (Figs. S5B&C), which is well-established as a non-covalent, weak inhibitor of SDH^33^. Altogether, these results suggest that Pam3 or Poly I:C alone do not elicit ‘breaks’ in the TCA cycle after 24 hr., and engaging both MyD88- and TRIF-linked pathways is required.

**Figure 4:**
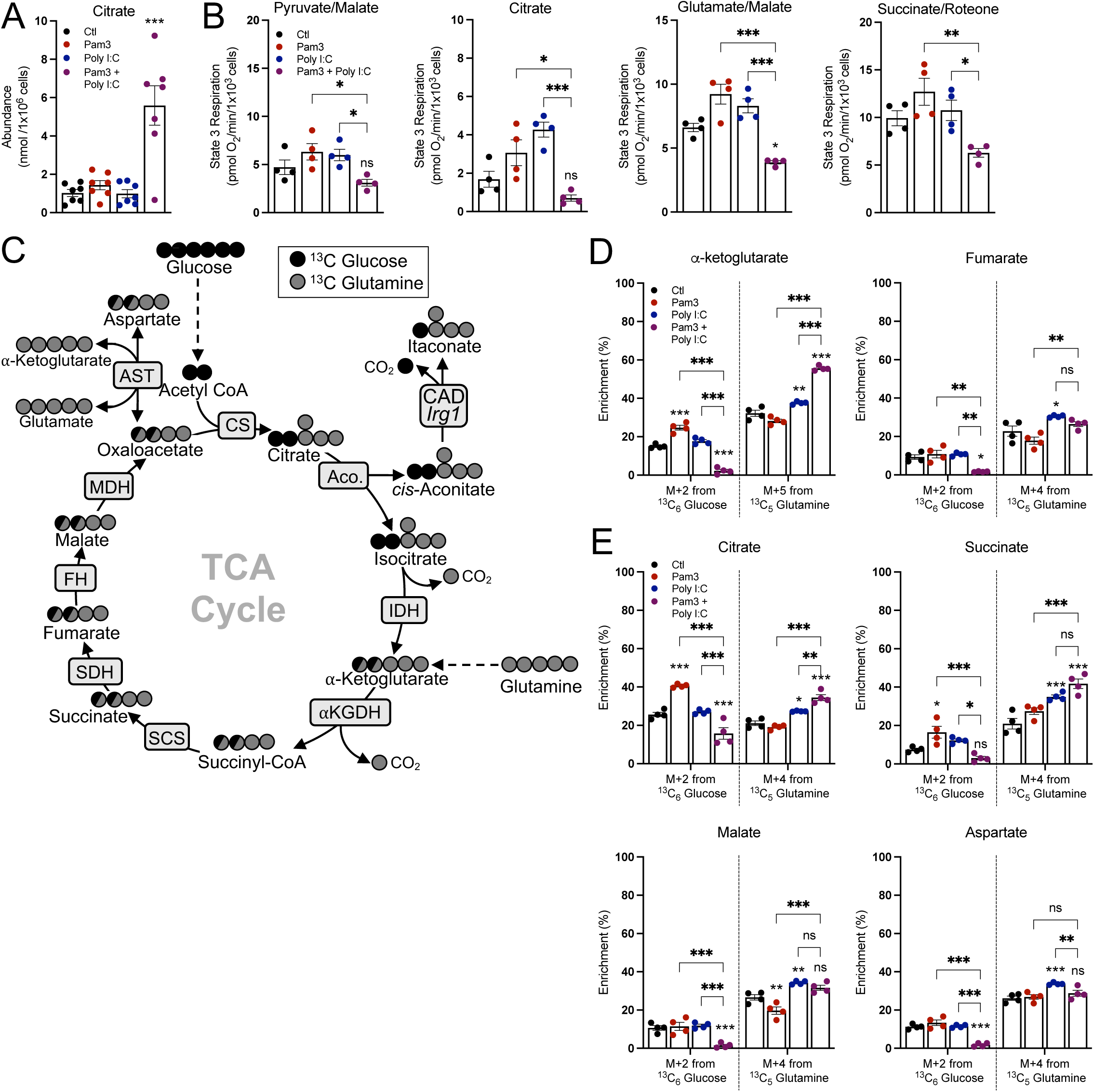
Neither Pam3 nor Poly I:C induce characteristic breaks in the TCA cycle. **A**) Intracellular abundance of citrate from control (Ctl) and BMDMs treated with Pam3, Poly I:C, or Pam3 + Poly I:C for 24 hr. (n = 7). **B**) State 3 respiration from permeabilized control (Ctl) and BMDMs treated with Pam3, Poly I:C, or Pam3 + Poly I:C for 24 hr. Permeabilized BMDMs were offered pyruvate/malate, citrate, glutamate/malate, or succinate/rotenone as substrates (n = 4). **C**) A graphical schematic depicting the labeling patterns of TCA cycle metabolites from ^13^C_6_ Glucose (black circles) and ^13^C_5_ Glutamine (grey circles). CS, citrate synthase; Aco., aconitase; IDH, isocitrate dehydrogenase; α-KGDH, alpha-ketoglutarate dehydrogenase; SCS, succinyl CoA synthetase; SDH, succinate dehydrogenase; FH, fumarate hydratase; MDH, malate dehydrogenase; AST, aspartate transaminase. **D&E**) Percent enrichment of denoted isotopologues from either ^13^C_6_ Glucose or ^13^C_5_ Glutamine from control (Ctl) and BMDMs treated with Pam3, Poly I:C, or Pam3 + Poly I:C for 24 hr. BMDMs were treated in unlabeled media for 16 hr. then changed to tracing media with stimuli for 6 hr. (n = 4). All data are mean ± SEM with statistical analysis conducted on data from biological replicates, each of which included multiple technical replicates, unless otherwise indicated. Statistical analysis for (**A**) and (**B**) was performed as an ordinary one-way, ANOVA followed by Tukey’s *post hoc* multiple comparisons test. Statistical analysis for (**D**) and (**E**) was performed as an ordinary two-way, ANOVA followed by Tukey’s *post hoc* multiple comparisons test.

We then hypothesized that Pam3- or Poly I:C-treated BMDMs accumulate signaling metabolites not by slowing metabolite consumption but rather by increasing synthesis. To test this hypothesis, we conducted stable isotope tracing in BMDMs offered ^13^C_6_-glucose and ^13^C_5_-glutamine to better understand TCA cycle metabolism in response to these stimuli. BMDMs were pretreated with TLR ligands for 18 hr., after which time cells were washed and given new medium containing ^13^C_6_-glucose or ^13^C_5_-glutamine for 6 hr. Tracing with isotopically labeled glucose or glutamine resulted in distinct labeling patterns, with glucose enriching the M+2 isotopologues of TCA cycle metabolites and glutamine enriching the M+5 or M+4 isotopologues (Fig. 4C). BMDMs treated with Pam3 showed increased enrichment from glucose into several TCA cycle metabolites, while this was predictably decreased in BMDMs co-treated with both Pam3 and Poly I:C in a manner consistent with reduced IDH activity (Fig. 4D, full isotopologue distributions from both ^13^C_6_-glucose and ^13^C_6_-glutamine can be found in Supplementary Table 1). Overall, in contrast to the co-treated BMDMs, macrophages treated with either Pam3 or Poly I:C for 24 hr. maintain relative TCA cycle flux and accumulate signaling metabolites with a different balance between metabolite synthesis and consumption.

### Respiratory inhibition does not enhance pro-inflammatory macrophage activation

After determining that accumulation of itaconate and succinate is independent of decreased oxidative phosphorylation, we next asked what other features of the macrophage pro-inflammatory response might be driven by respiratory inhibition. Indeed, a priming effect from IFN-γ on LPS-induced *Nos2* gene expression and nitric oxide (NO) production has been appreciated for decades^36,37^. We therefore hypothesized that combinations of MyD88- and IFN-linked ligands could amplify pro-inflammatory gene expression. As expected, Pam3 and Poly I:C synergistically increased the expression of many pro-inflammatory genes (Fig. 5A, S6A) after 24 hr., as did IMQ with Poly I:C (Figs. S6B), and 10 ng/mL LPS with IFN-γ (Figs. S6C). Co-treatment with Pam3 and Poly I:C also increased IL-1β and IL-12 release as well as NO production (Fig. 5B). These co-treatments also decreased mitochondrial respiration (Fig 1), establishing an association between enhanced pro-inflammatory gene expression and decreased mitochondrial respiration after 24 hr. To determine if this relationship was causative, we analyzed respiration and pro-inflammatory markers in BMDMs isolated from mice lacking inducible nitric oxide synthase (*Nos2,* iNOS). NO production via *Nos2* is the dominant mechanism by which oxidative phosphorylation decreases in macrophages^7–9,38,39^. As predicted, respiratory inhibition from co-treatment with Pam3 and Poly I:C was almost entirely rescued upon loss of iNOS (Figs. 5C, S7A&B). However, in line with previous reports, expression of many pro-inflammatory genes and release of cytokines was not significantly reduced (Figs. 5D&E, Fig. S7C)^7,9^. Thus, preventing the NO-mediated inhibition of the mitochondrial respiratory chain and TCA cycle does not affect pro-inflammatory gene expression.

**Figure 5:**
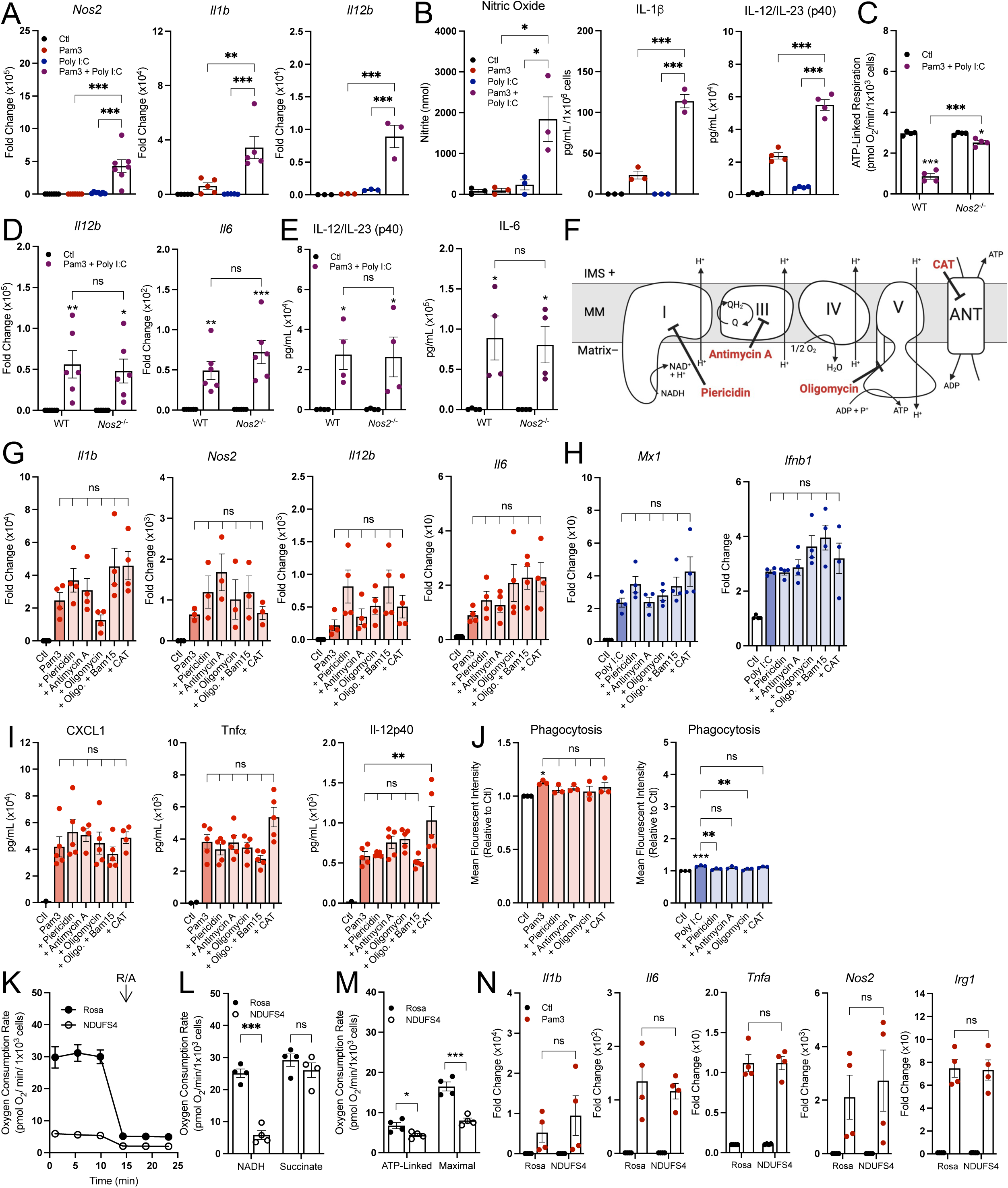
Respiratory inhibition does not enhance pro-inflammatory macrophage activation. **A**) Pro-inflammatory gene expression in control (Ctl) and BMDMs treated with Pam3, Poly I:C, or Pam3 + Poly I:C for 24 hr. relative to control (Ctl) (n = 3-7). **B**) Levels of nitric oxide (n = 3; not detected represented as zero), IL-1β (n = 3; not detected represented as zero), and IL-12/IL-23 (p40) (n = 4, values below the standard curve represented as zero) in medium collected from control (Ctl) and BMDMs treated with Pam3, Poly I:C, or Pam3 + Poly I:C for 24 hr. **C**) ATP-linked respiration rate for wildtype (WT) or iNOS-null (*Nos2*^-/-^) control (Ctl) and BMDMs treated with Pam3 + Poly I:C for 24 hr. (n = 4). **D&E**) Pro-inflammatory gene expression (n = 6) (**D**) and secreted cytokine levels of IL-12β from WT and Nos2^-/-^ control (Ctl) and BMDMs treated with Pam3 + Poly I:C for 24 hr. (n = 4; values below the standard curve represented as zero) (**E**). **G)** Schematic of mitochondrial respiratory chain with action of inhibitors used in G-J. CAT, carboxyatractyloside. **G&H**) Pro-inflammatory gene expression for control (Ctl) and BMDMs treated with Pam3 (n = 3-4) (**G**) or Poly I:C in combination with mitochondrial effector compounds (100 nM piericidin, 30 nM antimycin A, 10 nM oligomycin, 10 nM oligomycin + 3 µM Bam15, 30 µM CAT) for 24 hr. relative to control (Ctl) (n = 4) (**H**). **I**) Cytokine levels from medium of control (Ctl) and BMDMs treated with Pam3 in combination with mitochondrial effector compounds for 24 hr. (n = 5; values below the standard curve represented as zero). **J**) Phagocytosis of control (Ctl) and BMDMs treated with Pam3 or Poly I:C in combination with mitochondrial effector compounds for 24 hr. (n = 3). **K**) Representative oxygen consumption rates for Rosa and NDUFS4 knockdown B16 immortalized macrophage, O, oligomycin; F, FCCP; R/A, rotenone/antimycin A (n = 1 biological with 5 technical replicates) Where not visible, error bars are obscured by the symbol. **L&M**) State 3 respiration from B16 provided either NADH or succinate and (**L**) ATP-linked and maximal respiration for control (*Rosa*) and *Ndufs4* knockdown B16 (n = 4) (**M**). **N**) Pro-inflammatory gene expression in CRISPR edited B16 macrophages treated with Pam3 for 24 hr. relative to control (Ctl) (n = 4). All data are mean ± SEM with statistical analysis conducted on data from biological replicates, each of which included multiple technical replicates, unless otherwise indicated. Statistical analysis for (**A**), (**B**), and (**G-J**) was performed as an ordinary one-way, ANOVA followed by Tukey’s *post hoc* multiple comparisons test. Statistical analysis for (**C**), (**L**), and (**N**) was performed as an ordinary two-way, ANOVA followed by Sídák’s *post hoc* multiple comparisons test. Statistical analysis for (**D-E**) was performed as an ordinary two-way, ANOVA followed by Tukey’s *post hoc* multiple comparisons test. Statistical analysis for (**M**) was performed as an unpaired, two-tailed t-test.

To further understand whether reduced respiratory chain activity directly enhances classical macrophage activation, we measured whether inhibition of oxidative phosphorylation could enhance gene expression (Fig. 5F). When used in conjunction with a MyD88- or TRIF-linked TLR agonist for 24 hr., inhibition of complex I (with piericidin A), complex III (with antimycin A), the adenine nucleotide translocase (with carboxyatractyloside), or complex V (with oligomycin) did not enhance pro-inflammatory gene expression or cytokine release above that of Pam3, Poly I:C, or 10 ng/mL LPS alone (Figs. 5G-I, S8A-C). Additionally, the mitochondrial effector compounds had no effect on phagocytosis in combination with Pam3, and some even induced a moderate defect in Poly I:C-driven phagocytosis (Fig. 5J). Importantly, the inhibitors were used at the lowest concentration that elicited a maximal effect on respiration, and none of the compounds decreased cell counts (Figs. S8D-F). Finally, we CRISPR-edited immortalized primary macrophages to lack *Ndufs4*, a subunit of mitochondrial complex I. Despite a profound reduction in NADH oxidation that manifested in reduced ATP-linked and maximal respiration in macrophages (Figs. 5K-M), we observed no change in Pam3-linked pro-inflammatory gene expression in *Ndufs4*-depleted macrophages (Fig. 5N). Together, these results indicate that the relationship between pro-inflammatory gene expression and mitochondrial respiration is largely associative rather than causative.

### The induction of the pro-inflammatory response does not temporally align with respiratory inhibition

Although we observed no causal relationship between respiratory inhibition and macrophage activation at 24 hr., this did not obviate a role for oxidative phosphorylation at an earlier timepoint and during the induction of the pro-inflammatory response. Upon TLR agonism, expression of many genes associated with pro-inflammatory activation peaks within minutes or hours, and recedes soon after to taper inflammation^40–42^. We found that across eleven different pro-inflammatory genes, expression increased sharply soon after treatment with 50 ng/mL LPS, peaking within 3-6 hr. (Figs. 6A, S9A). However, much of our understanding regarding the role of oxidative phosphorylation in the pro-inflammatory response is based on data in response to LPS treatment ± IFN-γ for 24 hr. or longer^5,7,8,14,15,43^. We therefore sought to understand whether mitochondrial energetics were altered within a timeframe commensurate with the peak expression of canonical pro-inflammatory genes.

**Figure 6:**
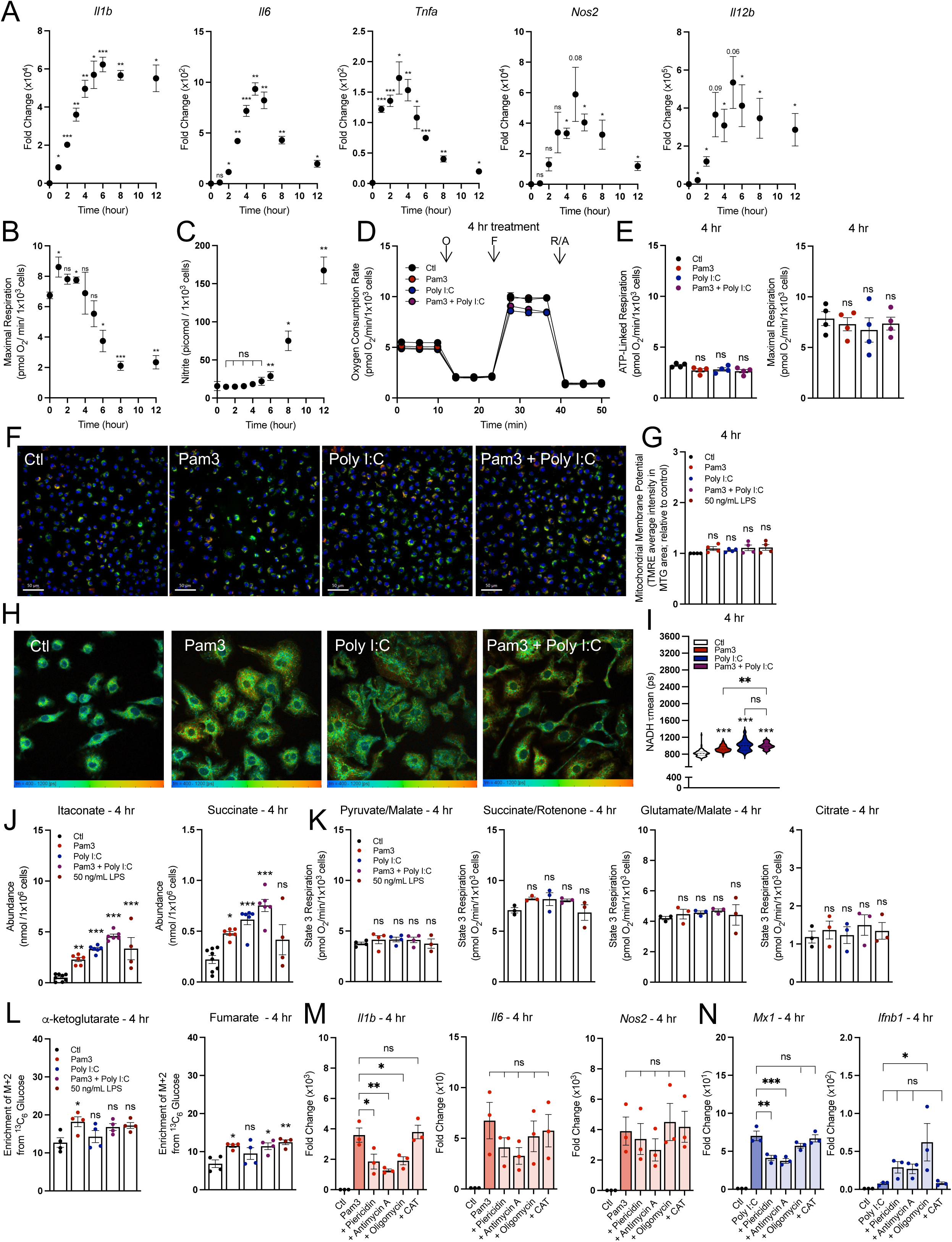
The induction of the pro-inflammatory response does not temporally align with respiratory inhibition. **A**) Pro-inflammatory gene expression in control (Ctl) and BMDMs treated with 50 ng/mL LPS across multiple timepoints relative to control (Ctl) (n = 3-4). **B**) Maximal respiration rate for control (Ctl) and BMDMs treated with 50 ng/mL LPS across multiple timepoints (n = 3-4). **C**) Levels of nitric oxide in medium from control (Ctl) and BMDMs treated with 50 ng/mL LPS across multiple timepoints (n = 3-4; not detected represented as zero). **D**) The oxygen consumption rates from a representative experiment with control BMDMs (Ctl) and BMDMs activated with Pam3, Poly I:C, or Pam3 + Poly I:C for 4 hr. Where not visible, error bars are obscured by the symbol. O, oligomycin; F, carbonyl cyanide-p-trifluoromethoxyphenylhydrazone (FCCP); R/A, rotenone/antimycin A (n = 1 biological with 5 technical replicates). **E**) ATP-Linked and maximal respiration rates for control (Ctl) and BMDMs treated with Pam3, Poly I:C, or Pam3 + P for 4 hr. (n = 4). **F**) Representative images of control (Ctl) and BMDMs treated with Pam3, Poly I:C, or Pam3 + Poly I:C for 4 hr. Nuclei are stained with Hoechst, mitochondria are stained with MTG and membrane potential as determined by TMRE. **G**) Bulk membrane potential as measured by Tetramethylrhodamine, ethyl ester (TMRE) fluorescence per mitochondrial area detected by MitoTracker Green (MTG) for control (Ctl) and BMDMs treated with Pam3, Poly I:C, or Pam3 + Poly I:C for 4 hr. Data is shown relative to control (n = 4). **H**) Representative images of control (Ctl) and BMDMs treated with Pam3, Poly I:C, or Pam3 + Poly I:C for 24 hr. analyzed via fluorescence lifetime imaging (FLIM). **I**) Mean endogenous NADH lifetime (**τ**mean) for whole cell measured in picoseconds (ps) from control (Ctl) and BMDMs treated with Pam3, Poly I:C, or Pam3 + Poly I:C for 4 hr. Data is from one biological replicate with each data point representing one cell (n = 87-135). **J**) Intracellular abundances of itaconate and succinate from control (Ctl) and BMDMs treated with Pam3, Poly I:C, or Pam3 + Poly I:C for 4 hr. (n = 4). **K**) State 3 respiration from permeabilized control (Ctl) and BMDMs treated with Pam3, Poly I:C, or Pam3 + Poly I:C for 4 hr. Permeabilized BMDMs were offered pyruvate/malate, citrate, glutamate/malate, or succinate/rotenone as substrates (n = 4). **L**) Percent enrichment of denoted isotopologues from ^13^C_6_ Glucose from control (Ctl) and BMDMs treated with Pam3, Poly I:C, or Pam3 + Poly I:C for 4 hr. (n = 4). **M&N**) Pro-inflammatory gene expression for control (Ctl) and BMDMs treated with Pam3 (n = 3) (**M**) or Poly I:C in combination with mitochondrial effector compounds (100 nM piericidin, 30 nM antimycin A, 10 nM oligomycin, 10 nM oligomycin + 3 µM Bam15, 30 µM CAT) for 4 hr. relative to control (Ctl) (n = 3) (**N**). All data are mean ± SEM with statistical analysis conducted on data from biological replicates, each of which included multiple technical replicates, unless otherwise indicated. Statistical analysis for (**A**) and (**C-D**) was performed as a paired, two-tailed t-test. Statistical analysis for (**F**), (**H**), and (**J-N)** was performed as an ordinary one-way, ANOVA followed by Tukey’s *post hoc* multiple comparisons test.

We measured rates of oxygen consumption at time-points between 1 and 12 hr. after treatment with 50 ng/mL LPS, and observed no significant defect in maximal respiration until 6 hr. and later (Fig. 6B). These initial findings suggest that even at an earlier timepoints, reduced oxidative phosphorylation does not regulate induction of pro-inflammatory gene expression given the lack of temporal alignment. As nitric oxide mediates the inhibition of mitochondrial respiration, we also measured nitrite levels (a stable product of nitric oxide degradation) at each timepoint (Fig. 6C). Indeed, respiratory inhibition was more closely aligned with the timeframe of nitrite accumulation rather than gene expression.

Having established that decreased mitochondrial respiration does not align temporally with the induction of pro-inflammatory gene expression, we measured other features of mitochondrial function during an early timeframe. Specifically, we determined whether co-treatment with Pam3 and Poly I:C displays the same profound mitochondrial alterations at 4 hr. (when gene expression is closer to its peak) as at 24 hr. However, we observed no alterations to mitochondrial respiration with either ligand alone or in combination after 4 hr. (Figs. 6D&E). We also observed no change in membrane potential for all groups relative to vehicle controls (Figs. 6F&G). We did, however, observe an increase in NADH fluorescence lifetime for all treatment groups relative to control, suggesting that the NADH pool size or redox status can be altered during macrophage activation while maintaining rates of oxidative phosphorylation (Figs. 6H&I).

We then measured whether steady-state metabolite levels could be increased after only 4 hr. treatment and prior to reductions in oxygen consumption. Indeed, itaconate and succinate levels increased at this relatively early time point for each treatment group, further suggesting that metabolite accumulation is independent of respiratory inhibition (Fig. 6J). We also observed no defects in permeabilized cell respirometry after 4 hr. treatment (Fig. 6K). Furthermore, unlike the 24 hr. timepoint (Fig. 4D), BMDMs co-treated with Pam3 and Poly I:C did not have decreased incorporation of ^13^C_6_-glucose into either α-ketoglutarate or fumarate, the two metabolites immediately downstream of the canonical ‘breaks’ in the TCA cycle (Fig. 6L).

Lastly, we treated examined whether pharmacologic inhibition of oxidative phosphorylation could increase gene expression stimulated by 4 hr. treatment of Pam3, Poly I:C, or LPS. Consistent with other data, we observed no enhancement in pro-inflammatory gene expression (Figs. 6M&N, S9B-D). In total, our data suggest reductions in mitochondrial respiration are not essential for the induction of the macrophage pro-inflammatory gene expression.

### Peritoneal macrophages can be activated in vivo without decreasing mitochondrial respiration

Finally, we assessed the effect of *in vivo* macrophage activation on mitochondrial energetics. Macrophages were intraperitoneally activated with Pam3 or LPS for 24 hours and assessed *ex vivo* 3 hr. after harvest (Figs. 7A; S10). Both treatments increased serum IL-6 as well as IL-6 and IL-12 in the lavage fluid (Figs. 7B, S10A) and LPS treatment resulted in increased pro-inflammatory gene expression (Fig. 7C), indicating macrophage activation. In *ex vivo* peritoneal macrophages, lactate efflux and intracellular lactate accumulation increased upon activation, again linking glycolysis with classical macrophage activation (Figs. 7D&E, S10C&D). Simultaneously, however, oxygen consumption rates were unchanged (Figs. 7E&F, S10E&F), further separating mitochondrial energetics from the induction of a pro-inflammatory response. Additionally, *ex vivo* peritoneal macrophages increased intracellular accumulation of itaconate and succinate but not citrate or α-ketoglutarate, showing again that pro-inflammatory metabolites can accumulate without conventional ‘breaks’ in the TCA cycle (Figs. 7G, S10E). To validate the model system, we activated peritoneal macrophages for 24 hr. *in vitro* and observed respiratory inhibition in response to LPS but not Pam3, showing that peritoneal macrophages behave similarly to BMDMs *in vitro* (Fig. S10F). Altogether, our work provides several independent lines of evidence that reductions in oxidative phosphorylation are not essential for the macrophage pro-inflammatory response.

**Figure 7:**
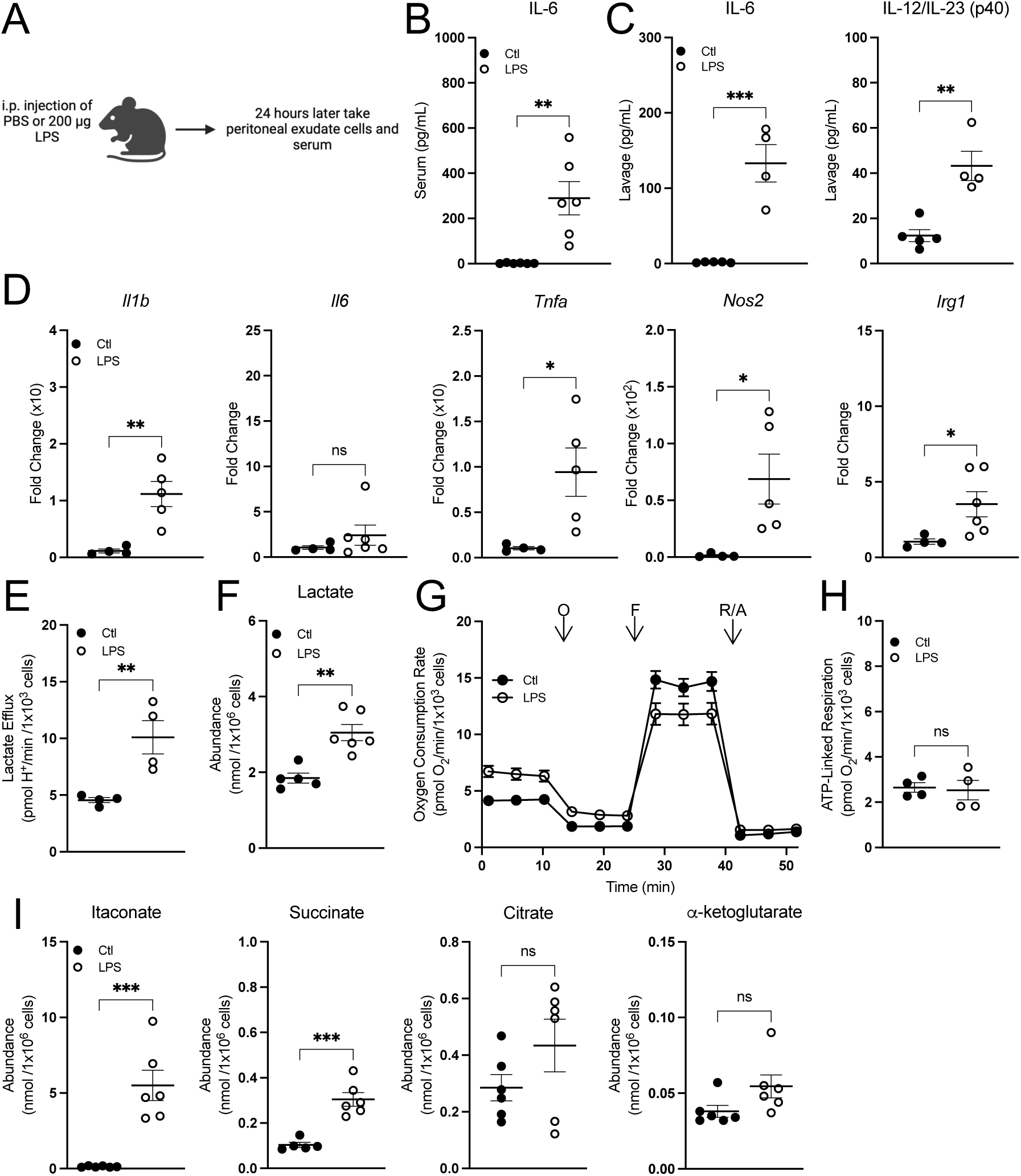
Peritoneal macrophages activated *in vivo* preserve oxidative phosphorylation and accumulate pro-inflammatory metabolites. **A**) A schematic depicting the experimental design for peritoneal macrophages. **B&C**) Cytokine levels from serum (n = 5-6) (**B**) and lavage fluid from mice intraperitoneally injected with PBS or LPS for 24 hr. (n = 4-6) (**C**). **D**) Pro-inflammatory gene expression from peritoneal macrophages isolated from mice intraperitoneally injected with PBS or LPS for 24 hr. relative to control (Ctl) (n = 4-6). **E**) Lactate efflux rates for peritoneal macrophages isolated from mice intraperitoneally injected with PBS or LPS for 24 hr. (n = 4). **F**) Intracellular lactate abundance from peritoneal macrophages isolated from mice intraperitoneally injected with PBS or LPS for 24 hr. (n = 5-6). **G**) Representative oxygen consumption trace with peritoneal macrophages isolated from mice intraperitoneally injected with PBS or LPS for 24 hr. Where not visible, error bars are obscured by the symbol. O, oligomycin; F, carbonyl cyanide-p-trifluoromethoxyphenylhydrazone (FCCP); R/A, rotenone/antimycin A (n = 1 biological replicate with 5 technical replicates). **H**) ATP-Linked respiration rates for peritoneal macrophages isolated from mice intraperitoneally injected with PBS or LPS for 24 hr. (n = 4). **I**) Intracellular itaconate, succinate, citrate and α-ketoglutarate abundances from peritoneal macrophages isolated from mice intraperitoneally injected with PBS or LPS for 24 hr. (n = 6). All data are mean ± SEM with statistical analysis conducted on data from biological replicates, each of which included multiple technical replicates, unless otherwise indicated. Statistical analysis for (**B-F**) and (**H-I**) was performed as an an unpaired, two-tailed t-test.

## DISCUSSION

Macrophage activation is often stratified as classical (Type I) pro-inflammatory activation or alternative (Type II) anti-inflammatory activation^44^, and in vitro studies of these activation states have shown these opposing functions are associated with equally polarized metabolic phenotypes^2^. However, as it is increasingly appreciated that macrophage polarization exists across a broad spectrum of activation states^45^, the metabolic phenotypes accompanying effector function also conceivably reside on a broad, flexible continuum^43,46,47^.

This work aligns with previous reports showing TLR agonism induces signal-specific metabolic reprogramming^48^. While both MyD88- and TRIF-dependent signaling pathways are required to alter mitochondrial respiration, the data suggest upregulation of glycolysis is largely driven by MyD88-dependent signaling. Additionally, our data indicate macrophages can accumulate succinate and itaconate without collapsing oxidative phosphorylation and ‘breaking’ the TCA cycle, but rather by rerouting TCA cycle metabolism to support both oxidative phosphorylation and synthesis of signaling metabolites. Interestingly, 24 hr. treatment with either Pam3 or Poly I:C resulted in different TCA cycle enrichment patterns from glucose or glutamine despite similar rates of oxidative phosphorylation, suggesting signal-specific regulation of intermediary metabolism.

The inability to amplify pro-inflammatory gene expression with respiratory chain inhibitors or CRISPR-mediated loss of the *Ndufs4* subunit of complex I contrasts results obtained with myeloid-specific loss of *Ndufs4*^17^. As such, further work is required to understand the temporal and developmental consequences of respiratory chain dysfunction. The work also highlights the importance of considering the duration, concentration, and specific stimuli used to elicit an *in vitro* pro-inflammatory response. Many of the hallmark mitochondrial alterations only occur under conditions of high nitric oxide production triggered by a combined MyD88 and IFN response. We and others have shown that human monocyte-derived macrophages that accumulate much smaller concentrations of nitric oxide do not display this characteristic mitochondrial repurposing^49^. Thus, some mitochondrial alterations observed in LPS-activated macrophages may be passive consequences of *in vitro* nitric oxide production and nitrite accumulation rather than requisite signals that drive macrophage function.

Of course, the results do not discount a role for other aspects of mitochondrial function in pro-inflammatory macrophage activation. For example, the increase in membrane potential observed with 24 hr. Pam3 treatment aligns with previous studies demonstrating that increasing mitochondrial membrane potential can enhance IL-1β production via RET-driven superoxide production^14^. Furthermore, even after only 4 hr. exposure to Pam3 or Poly I:C, macrophages exhibited a longer NADH fluorescence lifetime. This suggests redox changes could occur early in the pro-inflammatory response independently from alterations in oxidation phosphorylation or the mitochondrial membrane potential. As such, the work suggests that the generation of relevant mitochondrial signals – such as TCA cycle-linked metabolites and redox triggers – is entirely compatible with healthy oxidative phosphorylation and physiologically relevant bioenergetic parameters. Understanding precisely what these signals are will be essential to continue developing novel macrophage-targeted therapeutics^50^.

## ACKNOWLEDGEMENTS

ASD is supported by National Institutes of Health (NIH) Grants R35GM138003, the W.M. Keck Foundation (995337), and the Agilent Early Career Professor Award. ABB is supported by the UCLA Chemistry-Biology Interface Training Grant (T32GM136614). AEJ was supported by the UCLA Tumor Cell Biology Training Program (T32CA009056). SJB is supported by the NIH Grants P01HL146358 and R01HL157710. We also acknowledge the UCLA/CFAR Virology Core Lab (5P30AI028697), the UCLA Immune Assessment Core Facility, and the Bioenergetics and Imaging Core Facilities of the UCLA Metabolism Theme. Graphics in Figs. 1A, 1D, 3A, S5A, and 7A were created with BioRender.com.

## AUTHOR CONTRIBUTIONS

Andréa B Ball: Data curation; formal analysis; investigation; writing – original draft; writing – review and editing. Anthony E Jones: Data curation; formal analysis; investigation; methodology. Kaitlyn B Nguyễn: Data curation; formal analysis; investigation; writing – review and editing. Amy Rios: Data curation. Nico Marx: Data curation; formal analysis. Wei Yuan Hsieh: Data curation; formal analysis; investigation; methodology. Krista Yang: Data curation; formal analysis. Brandon R. Desousa: Data curation; formal analysis. Kristen K.O. Kim: Data curation; formal analysis. Michaela Veliova: Data curation; formal analysis. Zena Marie del Mundo: Data curation. Orian S. Shirihai: Conceptualization; resources. Cristiane Benincá: Data curation; formal analysis; methodology. Linsey Stiles: Data curation; formal analysis; resources; methodology. Steven J Bensinger: Conceptualization; resources; funding acquisition; methodology. Ajit S Divakaruni: Conceptualization; resources; data curation; formal analysis; supervision; funding acquisition; investigation; methodology; writing – original draft; project administration; writing – review and editing.

## METHODS

### Myd88^-/-^, Trif^-/-^, Ifnar^-/-^, Irg1^-/-^, and Nos2^-/-^ mice

Animal housing and all the experimental procedures were authorized by the UCLA Animal Research Committee. Mice were housed 4 per cage in a temperature (22°C-24°C) and humidity-controlled colony room, maintained on a 12 hr. light/dark cycle (07:00 to 19:00 light on), with a standard chow diet (LabDiet 5053) and water provided *ad libitum* with environmental enrichments. General health of the animal was assessed weekly by UCLA DLAM veterinarians.

The following strains were purchased from The Jackson Laboratory: C57BL/6J (strain # 000664); B6.129P2(SJL)-Myd88<tm1.1Defr>/J (strain # 009088); C57BL/6J-Ticam1<LPS2>/J (strain # 005037); C57BL/6NJ-Acod1/J (strain # 029340); B6.129P2-Nos2<TM1LAU>/J (strain # 002609). Femurs and tibias from *Ifnar*^-/-^ mice were generously provided by Dr. Ting-Ting Wu.

### Isolation of mouse bone marrow-derived macrophages (BMDMs)

Bone marrow cells were isolated from femurs of male mice between the age of 8-12 weeks as previously described^51^. Briefly, cells were treated with 3 mL RBC lysis buffer (Sigma-Aldrich) for 4 min to remove red blood cells, centrifuged at 400 *g* for 5 min, and resuspended in cell culture medium described below. Cells were maintained at 37°C in a humidified 5% CO_2_ incubator. BMDMs were differentiated for 6 days prior to experimental treatments, and medium was changed at day 4 of differentiation.

For all experiments involving BMDMs, cells were cultured in high-glucose DMEM (Gibco 11965) supplemented with 10% (v/v) heat-inactivated fetal bovine serum (FBS) unless otherwise indicated, 2 mM L-glutamine, 100 units/mL penicillin, 100 µg/mL streptomycin, 500 µM sodium pyruvate, and 5% or 10% v/v conditioned media containing macrophage colony stimulating factor (M-CSF) produced by CMG cells to induce differentiation to BMDMs^52^.

### Isolation of human PBMC-derived macrophages

Single donor human peripheral blood mononuclear cells (PBMC) were obtained from UCLA CFAR Virology Core lab for monocytes isolation. Monocytes were subsequently prepared by standard Ficoll isolation procedures and plastic adherence^53^. For macrophage differentiation, 1×10^7^ monocytes were suspended in DMEM supplemented with 10% (v/v) FBS with 2 mM L-glutamine, 100 units/mL penicillin, 100 µg/mL streptomycin, 500 µM sodium pyruvate, and 50 ng/mL of recombinant human M-CSF (Peprotech, 300-25) and cultured on non-tissue cultured treated Petri Dishes (Fisher, FB0875712) for 6 days prior to assays. Medium was changed at day 4 of differentiation.

### Isolation of mouse peritoneal macrophages

Mice were intraperitoneally injected with PBS, 200 µg LPS, or 200 µg Pam3 24 hr. prior to isolation of peritoneal macrophages. Mice were euthanized with isoflurane followed by cervical dislocation and abdominal skin was retracted to expose the intact peritoneal wall. 5 mL of ice-cold PBS with 2 mM EDTA and 2% FCS (Biochrom) was injected into the peritoneal cavity using a syringe with a 20-G needle. Following gentle massages to the cavity, the fluid was then aspirated from the peritoneal cavity using the same syringe and collected in a 15 mL tube. The procedure was repeated twice to obtain a final volume of 10 mL. The cell suspension was centrifuged at 400*g* for 5 min. Cell pellets were resuspended in BMDM culture medium and plated in 12-well plates. Cells were separated for 3 hr. at 37°C in a humidified 5% CO_2_ incubator with peritoneal macrophages adhering to the plate and other cells remaining in the supernatant. For peritoneal macrophages isolated for *in vitro* studies, mice were intraperitoneally injected with 3 mL of sterile thioglycolate broth for 72 hr. prior to isolation.

### Stimulation of bone marrow-derived macrophages

On day 6 after harvest, BMDMs were plated at different densities per well for the respective assays (see below). On day 8, macrophages were treated with 50 ng/mL LPS (or 10 ng/mL as noted in the figure legend), 50 ng/mL Pam3CSK4, 1 µg/mL Poly I:C, 10 µM imiquimod, 20 ng/mL IFN-γ, 20 ng/mL IFN-β, 100 nM CL307, 100 nM ODN1668 or simultaneously co-stimulated with a combination of the above for 24 hr. with controls. For the 4 hr. treatments, on day 8, the media was changed, and on day 9, macrophages were stimulated as noted in the figure legends.

### Mitochondrial effector compound treatment of BMDMs

For experiments involving respiratory chain inhibitors, BMDMs were vehicle-treated or treated with 100 nM piericidin A (Piericidin) (Sigma), 30 nM antimycin A (Sigma), 10 nM oligomycin (Sigma), 10 nM Oligomycin + 3 µM Bam15 (Sigma), or 30 µM carboxyatractyloside (CAT) (Sigma) for 24 hours. All inhibitors were given as co-treatments simultaneously with 50 ng/mL Pam3CSK4, 1 µg/mL Poly I:C, or 10 ng/mL LPS.

#### *In vitro* peritoneal macrophage stimulation

Freshly isolated thioglycolate-elicited peritoneal macrophages were seeded in Seahorse XF96 wells at a seeding density of 5 x 10^4^ cells/well in BMDM culture medium. The next day, cells were treated with either 50 ng/mL LPS or 50 ng/mL Pam3, and respiration was assessed after 24 hr.

### Seahorse XF Analysis

All respirometry was conducted in a Seahorse XF96 or XFe96 Analyzer (Agilent). All experiments were conducted at 37°C and at pH 7.4 (intact cells) or 7.2 (permeabilized cells). Respiration was measured in medium containing 8 mM glucose, 2 mM glutamine, 2 mM pyruvate, and 5 mM HEPES. Cells were plated at 5 x 10^4^ cells/well on day 6 and assayed on day 9 after treatments as described in figure legends. Respiration was measured in response to oligomycin (1 µM), carbonyl cyanide-p-trifluoromethoxyphenylhydrazone (FCCP) (0.75 nM or 1.5 µM), and rotenone (0.2 µM) with antimycin A (1 µM).

#### Intact Cells

Calculation of respiratory parameters were made according to standard protocols^54,55^. Briefly, ATP-linked respiration was calculated by subtracting the oxygen consumption rate insensitive to rotenone and antimycin A from the measurements after injection of oligomycin. Maximal respiration was calculated by subtracting the oxygen consumption rate insensitive to rotenone and antimycin A from the maximum rate obtained after injection of FCCP. Lactate efflux rates were calculated as previously described^56^.

For *ex vivo* peritoneal macrophages, cells were plated at 2.5 x 10^5^ cells/well in Cell-TAK (Corning) coated 96-well Seahorse XFe96 plates. Plates were spun at 500*g* for 4 min and respiratory parameters were obtained as previously described.

#### Permeabilized Cells

Recombinant, mutant perfringolysin O (rPFO; commercially XF Plasma Membrane Permeabilizer [XF PMP, Agilent Technologies]) was used to selectively permeabilize the plasma membrane of BMDMs. Experiments were conducted as previously described^57,58^. Immediately prior to assay, cell media was replaced with MAS buffer (70 mM sucrose, 220 mM mannitol, 10 mM KH_2_PO_4_, 5 mM MgCl_2_, 2 mM Hepes, and 1 mM EGTA; pH 7.2) containing 3 nM rPFO, respiratory substrates, and 4 mM ADP. The ADP-stimulated respiration rate (referred to as ‘State 3’ respiration) was measured, and rates were subsequently measured in response to 0.2 μM rotenone with 1 μM antimycin A. Substrate concentrations were as follows: Glutamate/Malate, 5 mM glutamate with 5 mM malate; Pyruvate/Malate, 5 mM pyruvate with 1 mM malate; Succinate/Rotenone, 5 mM succinate with 2 μM rotenone; Citrate, 5 mM citrate.

When permeabilized cells were treated with alamethicin to form pores of 3-6 kDa in the mitochondrial inner membrane (“double-permeabilized” cells) and complex I-mediated respiration was directly assessed, 10 μg/mL alamethicin was added at 37°C 15 minutes prior to measurements^58^. Double-permeabilized cells were offered 10 μM cytochrome c in the experimental medium and either 10 mM NADH or 10 mM succinate with 2 μM rotenone to drive respiration.

### Mitochondrial Membrane Potential

Cells were plated in the inner 60 wells of black-walled, 96-well plates at 30,000 cells/well. Prior to measurements of membrane potential, the medium was changed to high-glucose DMEM lacking serum and antibiotic but supplemented with 10 nM TMRE, 200 nM MitoTracker Green (MTG), and 1 µg/mL Hoechst. Cells were incubated in this medium for 1 hr. at 37°C. After incubation, cells were washed two times with this incubation medium lacking dye. Images were acquired with the 50 mm slit confocal mode and a 40x (1.2 NA) water lens in Z-stack mode of 0.5 mm slices with a total of 6 slices. Analysis was performed in the MetaXpress software keeping the same parameters for all the images acquired. Maximum Z-projections of MTG were used for morphologic analysis and the sum of Z-projections of TMRE was used for quantification of intensity. A TopHat filter was applied to the MTG images for better definition of structures and equalization of fluorescence. The images were thresholded and transformed into a binary segmentation. This segmented area was used to measure the average intensity of TMRE on pixels positive for MTG.

### Imaging studies

Live cell imaging of BMDMs was conducted on a Zeiss LSM880 using a 63x Plan-Apochromat oil-immersion lens and AiryScan super-resolution detector with a humidified 5% CO_2_ chamber on a temperature-controlled stage at 37°C. Cells were differentiated in glass-bottom confocal plates (Greiner Bio-One). BMDMs were incubated with 15 nM TMRE for 1 hr. in their regular culture medium. Image Analysis was conducted using FIJI (ImageJ, NIH). Image contrast and brightness were not altered in any quantitative image analysis protocols. Brightness and contrast were equivalently modified across measurement groups to allow proper representative visualization of the effects revealed by unbiased quantitation.

### Metabolite Quantification

Experiments were performed as previously described^59^. Briefly, BMDMs were plated at 1 x 10^6^ cells/well in 6-well plates and treated with macrophage stimuli as described above. Peritoneal macrophages were extracted immediately following the 3 hr. incubation in 12-well plates. Metabolite extraction was conducted with a Folch-like extraction with a 5:2:5 ratio of methanol:water:chloroform. 6- or 12-well dishes were kept on ice and quickly washed with ice-cold 0.9% (w/v) NaCl. Cells were then scraped in ice-cold methanol and water containing 5 µg/mL norvaline (Sigma #N7502), an internal standard. Chloroform was then added and samples were vortexed for 1 min and centrifuged at 10,000*g* for 5 min at 4°C.

The polar fraction (top layer) was removed, and the samples were dried overnight using a refrigerated CentriVap vacuum concentrator (LabConco). Metabolite standards (50 nmol to 23 pmol) were extracted alongside the cell samples to ensure the signal fell within the linear detection range of the instrument. The dried polar metabolites were reconstituted in 20 µL of 2% (w/v) methoxyamine in pyridine prior to a 45-min incubation at 37°C. Subsequently, 20 µL of MTBSTFA with 1% tert-butyldimethylchlorosilane was added to samples, followed by an additional 45-min incubation at 37°C. Samples were analyzed using Agilent MassHunter software. Samples were analyzed using a DB-35 column (Agilent Technologies). Information regarding additional technical specifications is available elsewhere^60,61^.

### Stable isotope tracing

On day 6 cells were seeded 1 x 10^6^ cells/well in 6-well plates in BMDM culture medium. For 4 hr. assays on day 8, medium was changed and on day 9, cells were treated for 4 hr. with ligands as indicated in the figure legend in medium containing either 10 mM ^13^C_6_ glucose (Cambridge Isotope Laboratories) or 6 mM ^13^C_5_ glutamine (Cambridge Isotope Laboratories). For 24 hr. assays on day 8, cells were treated with ligands as indicated in the figure legend in culture medium. On day 9, 18 hours later, the medium was changed to culture medium with ligands and either 10 mM ^13^C_6_ glucose (Cambridge Isotope Laboratories) or 6 mM ^13^C_5_ glutamine (Cambridge Isotope Laboratories) for 6 hr. For medium containing each stable isotope labeled metabolite, the other respective metabolite was still present at the same concentration though unlabeled. After incubation in medium containing a stable isotope labeled metabolite, metabolites were extracted as described above. FluxFix software (http://fluxfix.science) was used to correct for the abundance of natural heavy isotopes against an in-house reference set of unlabeled metabolite standards^62^.

### Quantitative real-time RT-PCR (qPCR)

For BMDM experiments involving gene expression analysis, day 6 BMDMs were seeded 3 x 10^5^ cells/well in 12-well plates in BMDM culture medium. For 4 hr. treatments, on day 8, medium was changed, and on day 9 BMDMs were treated with ligands as indicated in the figure legend in culture medium supplemented with 5% (v/v) FBS. For 24 hr. treatments, BMDMs were treated on day 8 with ligands as indicated in the figure legend in culture medium supplemented with 5% (v/v) FBS. Peritoneal macrophages were lysed immediately following the 3 hr. incubation in 12-well plates. Cells were collected in Qiagen RNeasy Cell Lysis Buffer and RNA was extracted according to manufacturer’s protocol (Qiagen). cDNA was synthesized using 1,000 ng RNA per reaction with high-capacity cDNA reverse transcription kit (Applied Biosystems). KAPA SYBR Green qPCR Master Mix (2X) Kit (Applied Biosystems) and an Applied Biosystems QuantStudio 5 were used for quantitative RT-PCR using 0.5 µmol/L primers. Fold change related to the control group was calculated using 2^ΔΔCT^ method with *36b4* as the reference gene.

The primers were (forward and reverse, respectively) 5’-GCCCATCCTCTGTGACTCAT-3’ and 5’-AGGCCACAGGTATTTTGTCG-3’ for *Il1b*; 5’-AGTTGCCTTCTTGGGACTGA-3’ and 5’-TCCACGATTTCCCAGAGAAC-3’ for *Il6*; 5’-TGCCTATGTCTCAGCCTCTTC-3’ and 5’-GAGGCCATTTGGGAACTTCT-3’ for *Tnfa*; 5’-CACCTTGGAGTTCACCCAGT-3’ and 5’-ACCACTCGTACTTGGGATGC-3’ for *Nos2*; 5’-ATCGTTTTGCTGGTGTCTCC-3’ and 5’-GGAGTCCAGTCCACCTCTACA-3’ for *Il12b*; 5’-GCAACATGATGCTCAAGTCTG-3’ and 5’-TGCTCCTCCGAATGATACCA-3’ for *Irg1*; 5’-GACCATAGGGGTCTTGACCAA-3’ and 5’-AGACTTGCTCTTTCTGAAAAGCC-3’ for *Mx1*; 5’-GAGGCTCTTCAGAATGAGCAAA-3’ and 5’-CTCTGCGGTCAGTCTCTCT-3’ for *Mx2*; 5’-CAGCTCCAAGAAAGGACGAAC-3’ and 5’-GGCAGTGTAACTCTTCTGCAT-3’ for *Ifnb1*; 5’-AACATCCAGAACAACTGGCGG-3’ and 5’-GTCTGACGTCCCAGGGCA-3’ for *Isg20*; 5’-GGCCGATACAAAGCAGGAGAA-3’ and 5’-GGAGTTCATGGCACAACGGA-3’ for *Irf1*; 5’-TCCAGTTGATCCGCATAAGGT-3’ and 5’-CTTCCCTATTTTCCGTGGCTG-3’ for *Irf7*; 5’-CAGGGAAAATGGAAGTGGTG-3’ and 5’-CAGAGAGGTTCTCCCGACTG-3’ for *Ifi204;* 5’-CTGTGCCAGCTCAGAACACTG-3’ and 5’-TGATCAGCCCGAAGGAGAAG-3’ for *36b4*.

### Griess Assay

Nitric oxide was measured from BMDM supernatant 24 hr. after effector treatments. Briefly nitrite, a stable product of nitric oxide degradation, was measured by mixing 50 µL of culture supernatants with 50 µL Griess Reagent (Sigma), incubating in the dark for 15 min at room temperature, and measuring absorbance at 540 nm. Sodium nitrite was used as the standard curve for calculation of picomole of nitrite.

### Cytokine measurements

Enzyme-linked immunosorbent assays (ELISAs) were used to measure mouse IL-6 and IL-12b/IL-23 (p40) in BMDM medium (supernatant collected after centrifugation), mouse serum, or mouse lavage fluid according to manufacturer’s instructions (BioLegend). IL-1β levels in BMDM cell-culture supernatant was studied by Luminex’s xMAP® Immunoassay with the UCLA Immune Assessment Core Facility. CXCL1 (KC), TNF-α, and IL-12p40 levels in BMDM cell-culture supernatant was studied by BioLegend’s LEGENDplex MU M1 Macrophage Panel (8-plex) with V-bottom plate according to manufacturer’s instructions.

### Cell counts and normalization

When normalizing respirometry experiments and metabolite quantification to cell number, BMDMs were fixed immediately upon completion of the assay with 2% (w/v) formaldehyde for 20 min at room temperature and kept refrigerated between 1 and 14 days until assessment. The day prior to cell counting, cells were stained with Hoechst 33342 (Thermo Fisher) at 10 ng/mL overnight at room temperature. Cell counts were obtained using the Operetta High Content Imaging System (Perkin Elmer).

### Statistical Analysis

All statistical parameters, including the number of biological replicates (n), can be found in the figure legends. Each data point represents a biological replicate obtained from an individual mouse/human sample and is comprised from the average of multiple technical replicates. Statistical analyses were performed using Graph Pad Prism 10 software. Data are presented as the mean ± SEM. Individual pairwise comparisons were performed using two-tailed Student’s t-test. For experiments involving two or more factors, data were analyzed by one-way, repeated measures ANOVA followed by Tukey’s *post hoc* multiple comparisons tests. For other multiple values comparisons, data were analyzed by ordinary two-way, ANOVA followed by Tukey’s or Sídák’s *post hoc* multiple comparisons test when required. Data were assumed to follow a normal distribution (no tests were performed). Values denoted as follows were considered significant: *p < 0.05; **p <0.002; ***p<0.001.

