## Supplementary Figures 1-10 for "Pro-inflammatory macrophage activation does not require inhibition of mitochondrial respiration"

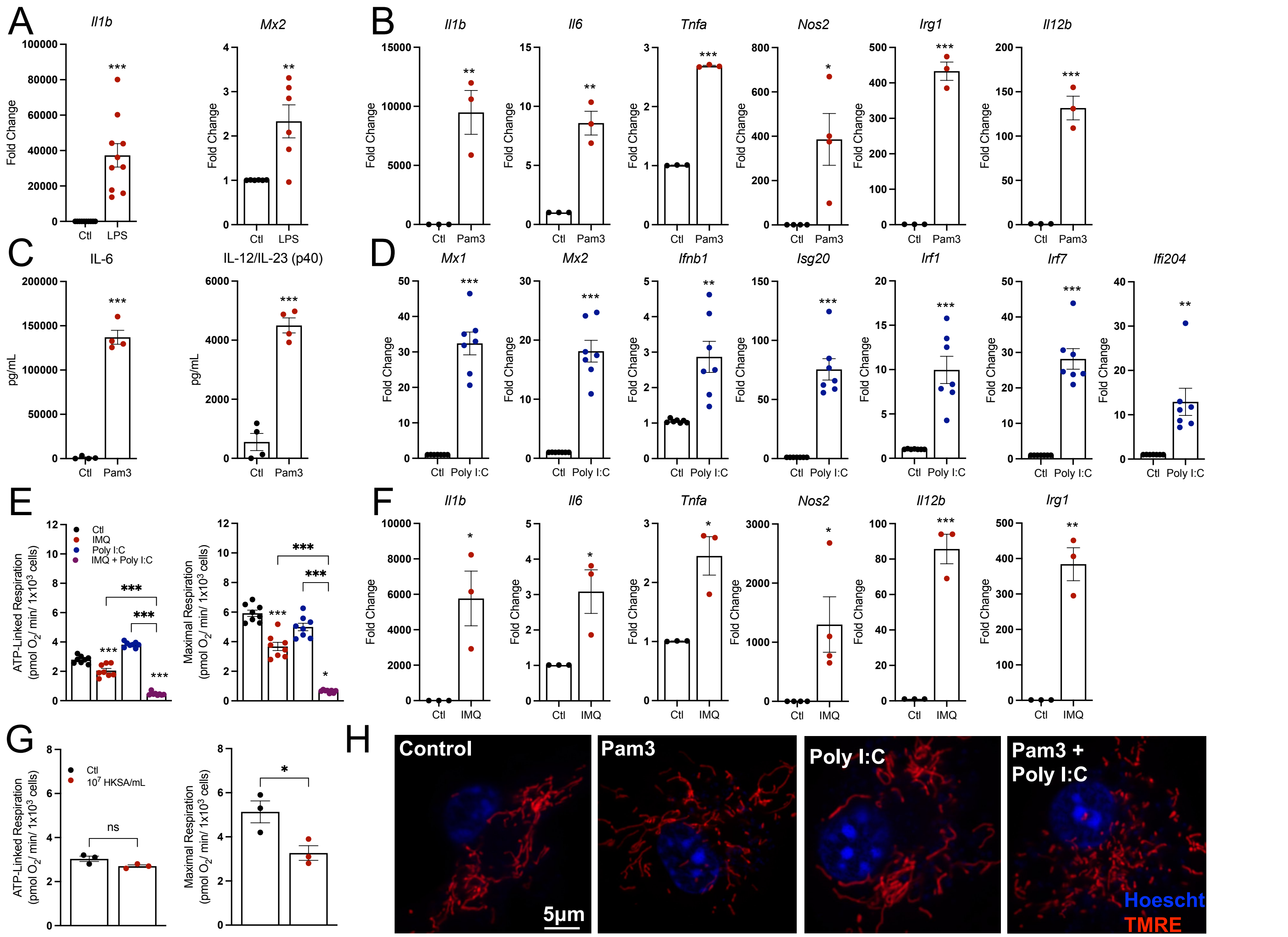


**Supplementary Figure 1: A**) Pro-inflammatory gene expression of control (Ctl) and BMDMs treated with 50 ng/mL LPS for 24 hr. (n = 6-10). **B**) Pro-inflammatory gene expression for BMDMs treated with Pam3 for 24 hr. (n = 3-4). **C**) Cytokine levels in medium from control (Ctl) and BMDMs treated with Pam3 for 24 hr. (n = 4; values below the standard curve are denoted as zero). **D**) Pro-inflammatory gene expression of control (Ctl) and BMDMs treated with Poly I:C for 24 hr. (n = 7). **E**) ATP-linked and maximal respiration rates for BMDMs treated with Imiquimod (IMQ), Poly I:C, or IMQ + Poly I:C (n = 8). **F**) Pro-inflammatory gene expression for BMDMs treated with IMQ for 24 hr. (n = 3-4). **G**) ATP-linked respiration, maximal respiration, and lactate efflux rates for control (Ctl) and BMDMs treated with $10^7$ heat-killed staphylococcus A (HKSA) for 24 hr. (n = 3). **H**) Representative images of mitochondrial morphology of BMDMs. Nuclei are stained with Hoechst and mitochondria are stained with TMRE. All data are mean ± SEM with statistical analysis conducted on data from biological replicates, each of which included multiple technical replicates, unless otherwise indicated. Statistical analysis for (**A-D**) and (**F-G**) was performed as an unpaired, two-tailed t-test**.** Statistical analysis for (**E**) and (**G**) was performed as an ordinary one-way, ANOVA followed by Tukey's *post hoc* multiple comparisons test.

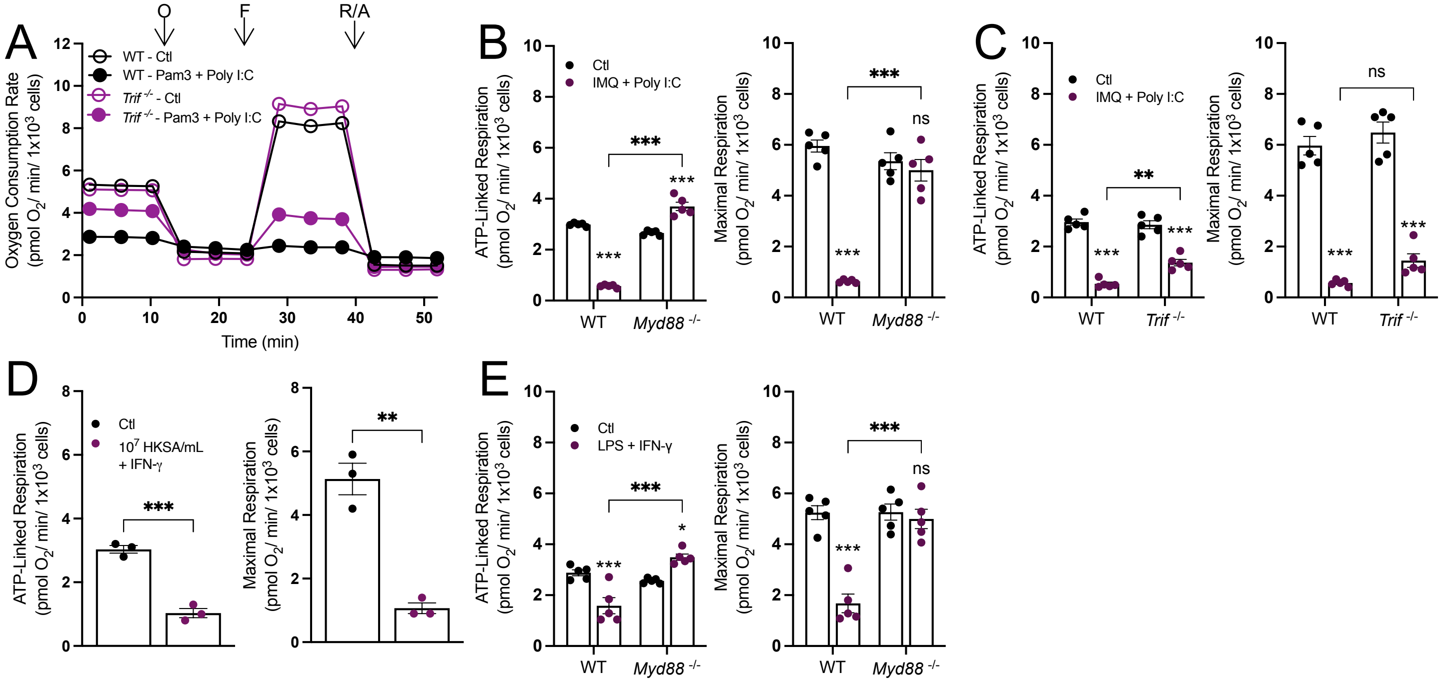


**Supplementary Figure 2: A**) The oxygen consumption rates from a representative experiment for wildtype (WT) and TRIF-null (*Trif*$^{-/-}$) control (Ctl) BMDMs or BMDMs treated with Pam3 + Poly I:C for 24 hr. O, oligomycin; F, FCCP; R/A, rotenone/antimycin A (n = 1 biological with 5 technical replicates). **B**) ATP-linked and maximal respiration calculations for WT and *Myd88*$^{-/-}$ control (Ctl) and BMDMs treated with imiquimod (IMQ) + Poly I:C (n = 5). **C**) ATP-linked and maximal respiration calculations for WT and *Trif*$^{-/-}$ control (Ctl) and BMDMs treated with IMQ + Poly I:C (n = 5). **D**) ATP-linked and maximal respiration calculations for WT control (Ctl) and BMDMs treated with $10^7$ heat-killed staphylococcus A (HKSA) + IFN-$\gamma$ (n = 3). **E**) ATP-linked and maximal respiration calculations for WT and *Myd88*$^{-/-}$ control (Ctl) and BMDMs treated with LPS + IFN-$\gamma$ (n = 5). All data are mean ± SEM with statistical analysis conducted on data from biological replicates, each of which included multiple technical replicates, unless otherwise indicated. Statistical analysis for (**B-E**) was performed as an ordinary two-way, ANOVA followed by Sídák's *post hoc* multiple comparisons test. Statistical analysis for (**F**) was performed as an unpaired, two-tailed t-test.

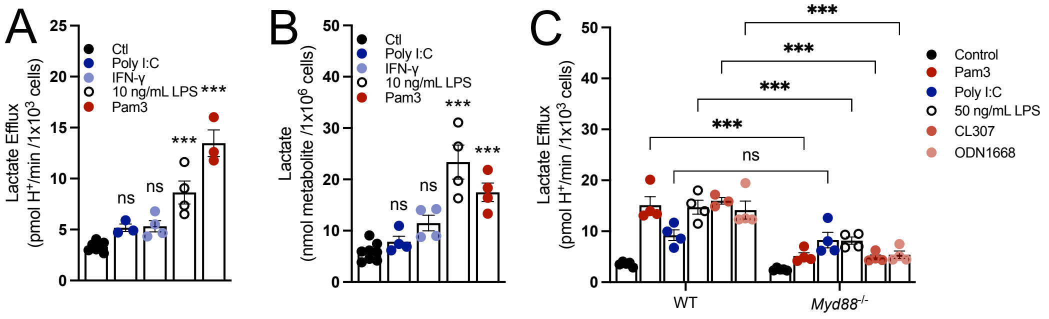


**Supplementary Figure 3: A**) Lactate efflux rates for control (Ctl) BMDMs or BMDMs treated with Poly I:C, IFN-gamma, 10 ng/mL LPS, or Pam3 for 24 hr. (n = 3-4). **B**) Intracellular lactate abundance for Ctl BMDMs or BMDMs treated with Poly I:C, IFN-gamma, 10 ng/mL LPS, or Pam3 for 24 hr. (n = 4). **C**) Lactate efflux rates from wildtype (WT) and MyD88-null (*Myd88*$^{-/-}$) control (Ctl) BMDMs or BMDMs treated with Pam3, Poly I:C, 50 ng/mL LPS, CL307, or ODN1668 for 24 hr. (n = 3-4). All data are mean ± SEM with statistical analysis conducted on data from biological replicates, each of which included multiple technical replicates, unless otherwise indicated. Statistical analysis for (**A**) and (**B**) was performed as an ordinary one-way, ANOVA followed by Tukey's *post hoc* multiple comparisons test. Statistical analysis for (C) was performed as an ordinary two-way ANOVA followed by Sídák's *post hoc* multiple comparisons test.

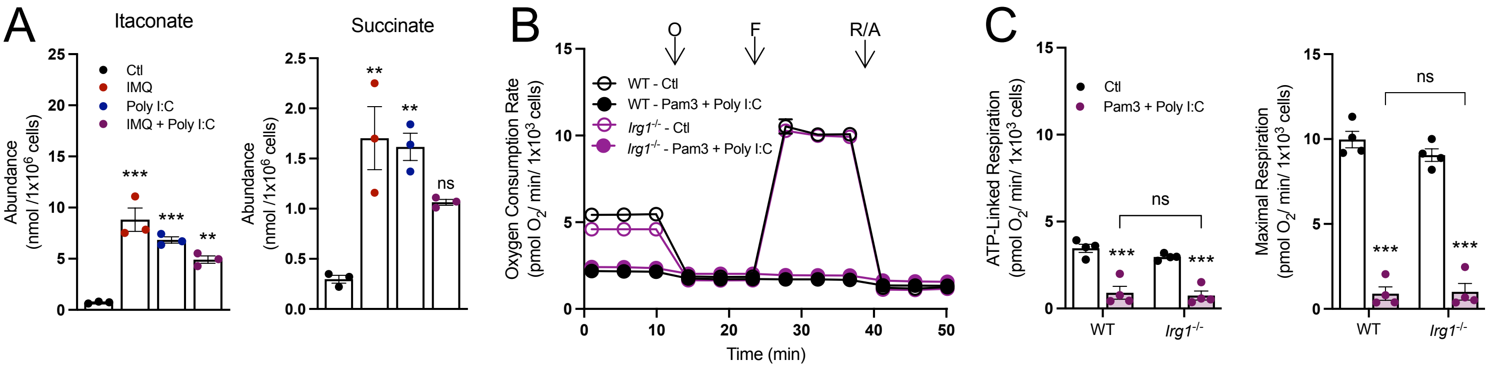


**Supplementary Figure 4: A**) Intracellular abundance of itaconate and succinate from control (Ctl) and BMDMs treated with IMQ, Poly I:C, or IMQ + Poly I:C (n = 3). **B**) The oxygen consumption rates from a representative experiment for wildtype (WT) and IRG1-null (*Irg1*$^{-/-}$) control (Ctl) and BMDMs activated with Pam3 + Poly I:C, O, oligomycin; F, carbonyl cyanide-p-trifluoromethoxyphenylhydrazone (FCCP); R/A, rotenone/antimycin A (n = 1 biological with 5 technical replicates). **C**) ATP-linked and maximal respiration rates for WT and *Irg1*$^{-/-}$ control (Ctl) and BMDMs treated with Pam3 + Poly I:C (n = 4). All data are mean ± SEM with statistical analysis conducted on data from biological replicates, each of which included multiple technical replicates, unless otherwise indicated. Statistical analysis for (**A**) was performed as an ordinary one-way, ANOVA followed by Tukey's *post hoc* multiple comparisons test. Statistical analysis for (**C**) was performed as an ordinary two-way, ANOVA followed by Sídák's *post hoc* multiple comparisons test.

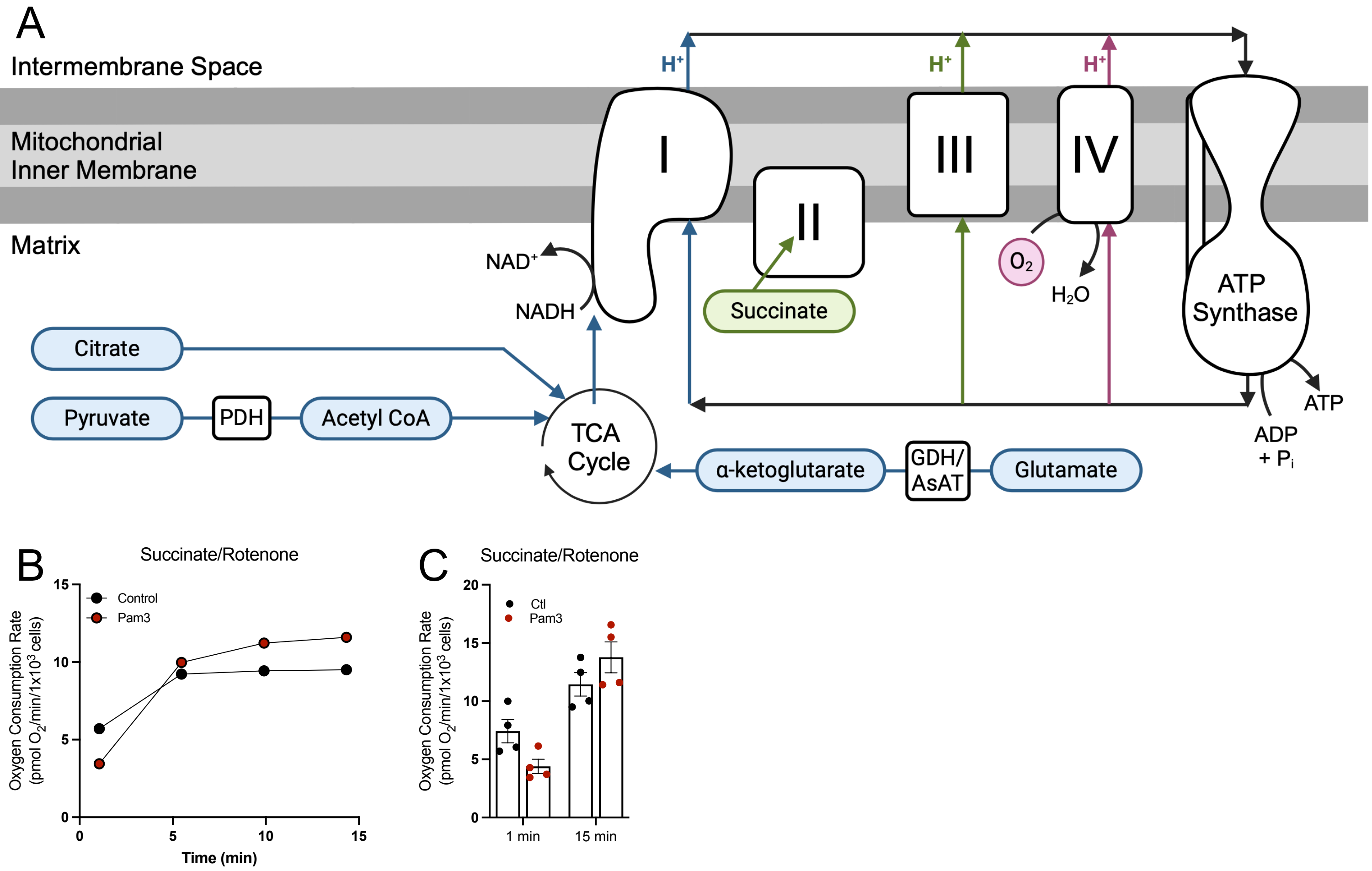


**Supplementary Figure 5: A**) A graphical schematic of permeabilized respirometry substrates feeding into the TCA cycle or electron transport chain. PDH, pyruvate dehydrogenase; GDH, glutamate dehydrogenase; AsAT, aspartate aminotransferase. **B**) The oxygen consumption rates from a representative permeabilized respirometry assay with succinate and rotenone as the substrates and control (Ctl) and BMDMs activated with Pam3 for 24 hr. O, oligomycin; F, carbonyl cyanide-p-trifluoromethoxyphenylhydrazone (FCCP); R/A, rotenone/antimycin A (n = 1 biological with 5 technical replicates). **C**) Rates of oxygen consumption after the first and fourth measurements for control (Ctl) and BMDMs treated with Pam3 for 24 hr., permeabilized, and provided succinate and rotenone as substrates (n = 4).

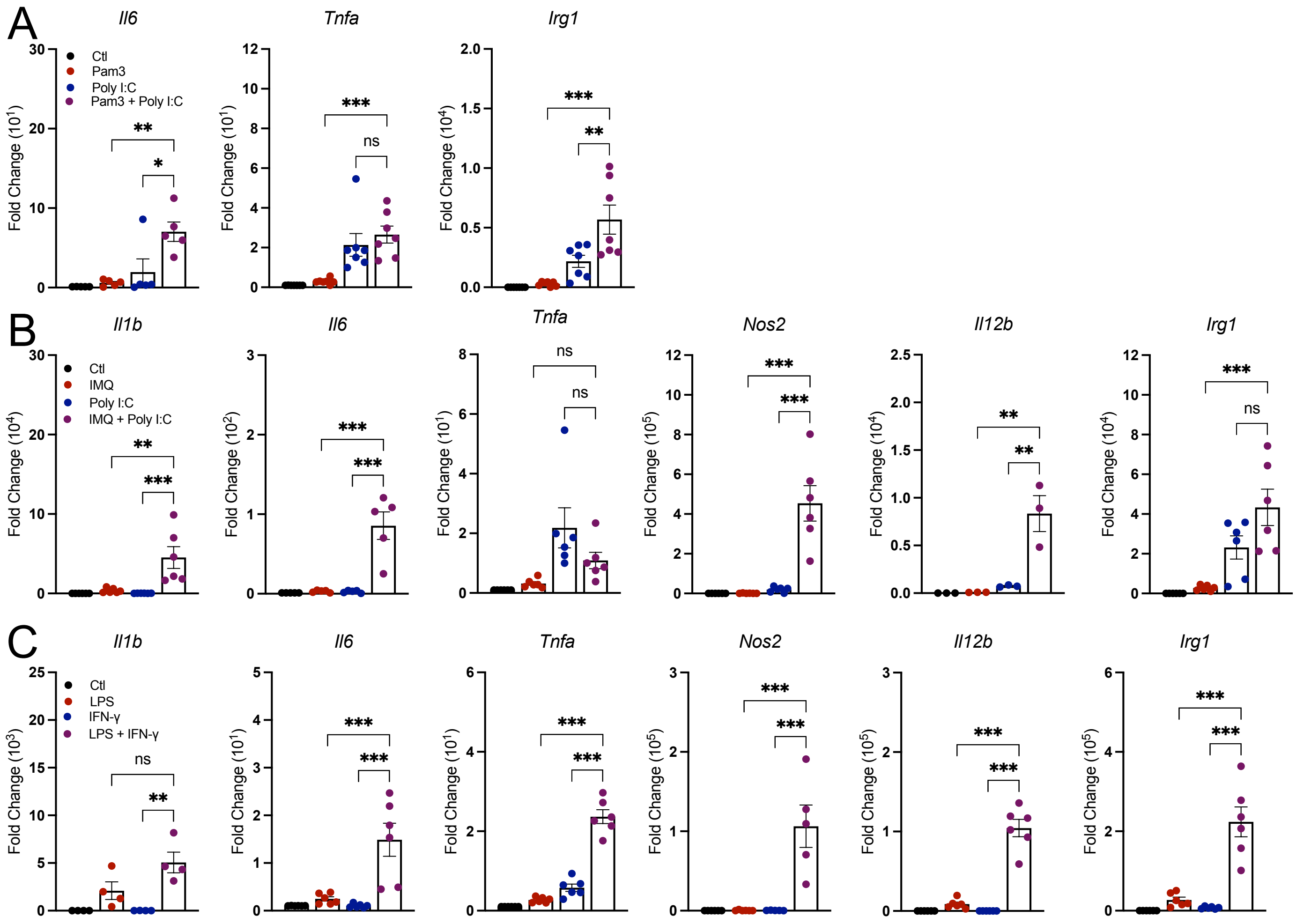


**Supplementary Figure 6: A-C**) Pro-inflammatory gene expression from BMDMs treated with the following groups: control (Ctl), Pam3, Poly I:C, or Pam3 + Poly I:C (n = 5-7) (**A**); control (Ctl), imiquimod (IMQ), Poly I:C, or IMQ + Poly I:C (n = 3-6) (**B**); or control (Ctl), 10 ng/mL LPS, IFN-$\gamma$, LPS + IFN-$\gamma$ (n = 4-6) (**C**). All data are mean ± SEM with statistical analysis conducted on data from biological replicates, each of which included multiple technical replicates, unless otherwise indicated. Statistical analysis for (**A-C**) was performed as an ordinary one-way, ANOVA followed by Tukey's *post hoc* multiple comparisons test.

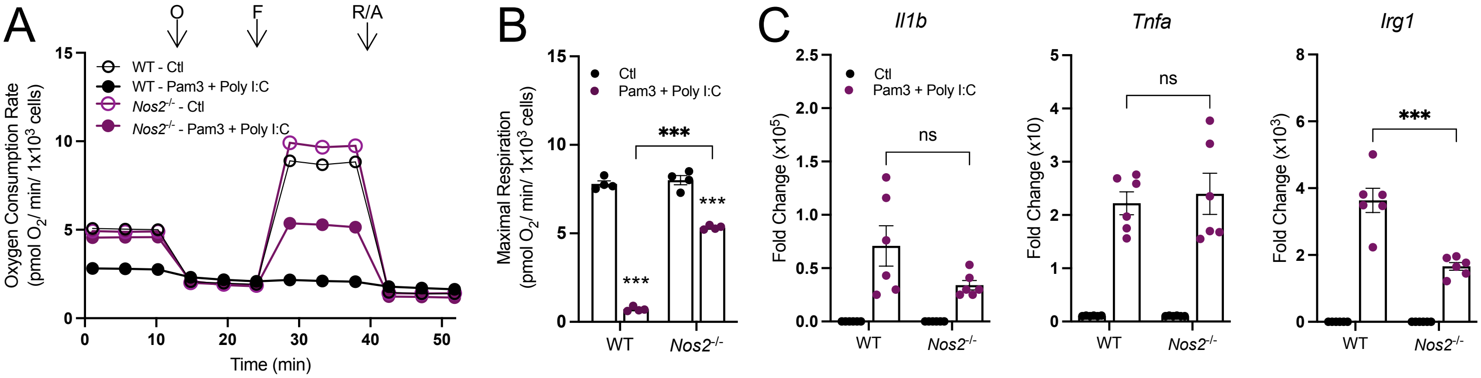


**Supplementary Figure 7: BMDMs from *Nos2*$^{-/-}$ mice have improved respiration but a comparable pro-inflammatory phenotype to WT BMDMs. A**) The oxygen consumption rates from a representative experiment for wildtype (WT) or iNOS-null (*Nos2*$^{-/-}$) control (Ctl) and BMDMs treated with Pam3 + Poly I:C., O, oligomycin; F, carbonyl cyanide-p-trifluoromethoxyphenylhydrazone (FCCP); R/A, rotenone/antimycin A (n = 1 biological with 5 technical replicates). **B**) Maximal respiration rates for WT and *Nos2*$^{-/-}$ control (Ctl) and BMDMs treated with Pam3 + Poly I:C (n = 4). **C**) Pro-inflammatory gene expression relative to control for WT and *Nos2*$^{-/-}$ control (Ctl) and BMDMs treated with Pam3 + Poly I:C for 24 hr. (n = 6). All data are mean ± SEM with statistical analysis conducted on data from biological replicates, each of which included multiple technical replicates, unless otherwise indicated. Statistical analysis for (**B**) and (**C**) was performed as an ordinary two-way, ANOVA followed by Tukey's *post hoc* multiple comparisons test.

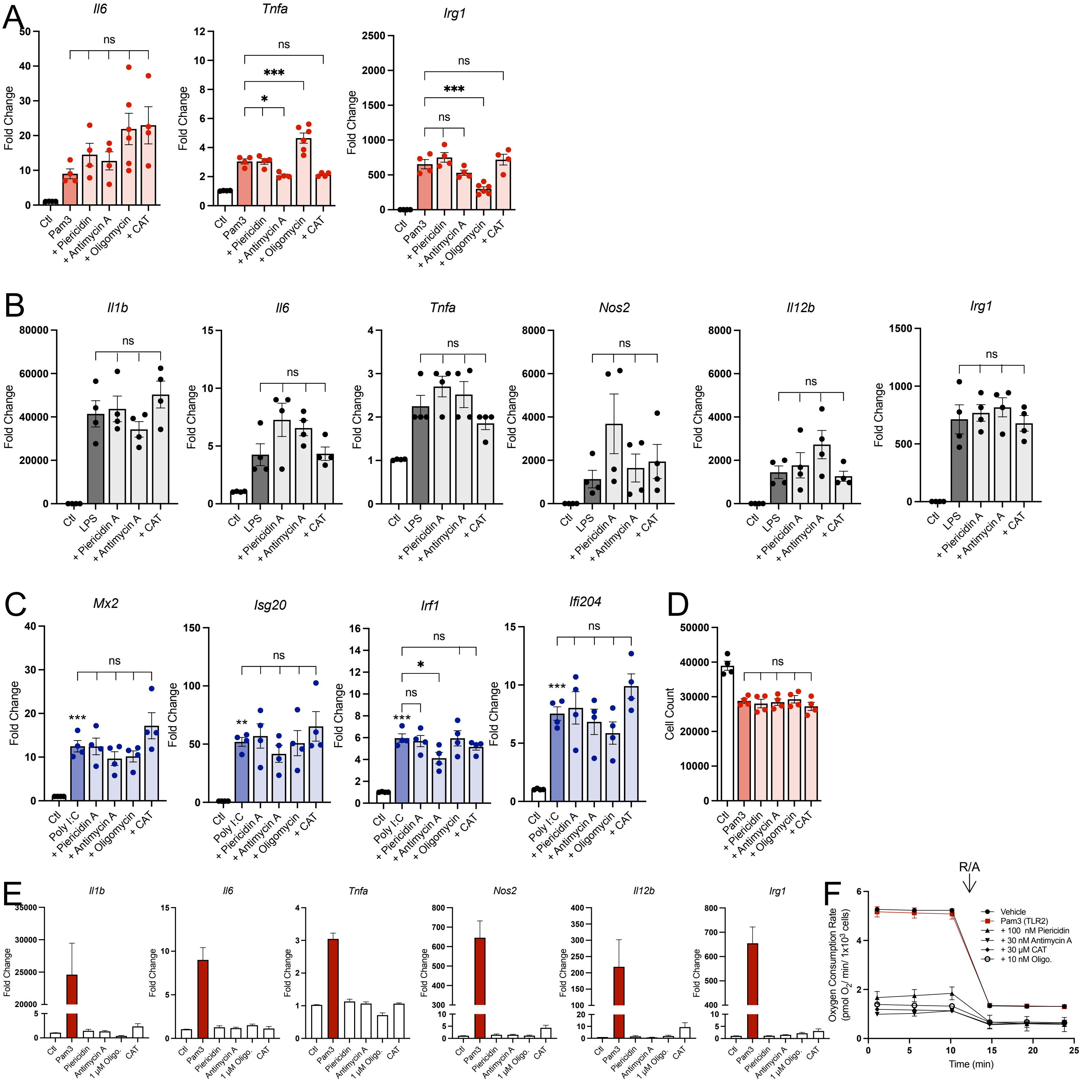


**Supplementary Figure 8: A-C**) Pro-inflammatory gene expression of control (Ctl) and BMDMs treated with Pam3 (n = 4-6) (**A**), LPS (n = 4) (**B**), or Poly I:C with mitochondrial effector compounds for 24 hr. (n = 4) (**C**). **D**) Cell counts via high-content imaging of Hoechst stained control (Ctl) and BMDMs treated with Pam3 with mitochondrial effector compounds for 24 hr. (n = 4). **E**) Pro-inflammatory gene expression of BMDMs treated with control (Ctl), Pam3 alone, or mitochondrial effector compounds alone for 24 hr. CAT, carboxyatractyloside (n = 3-4). **F**) Oxygen consumption rates for control (Ctl) and BMDMs treated with Pam3 with mitochondrial effector compounds for 24 hr. R/A, rotenone/antimycin A (n = 1 biological with 5 technical replicates). All data are mean ± SEM with statistical analysis conducted on data from biological replicates, each of which included multiple technical replicates, unless otherwise indicated. Statistical analysis for (**A-D**) was performed as an ordinary one-way, ANOVA followed by Tukey's *post hoc* multiple comparisons test.

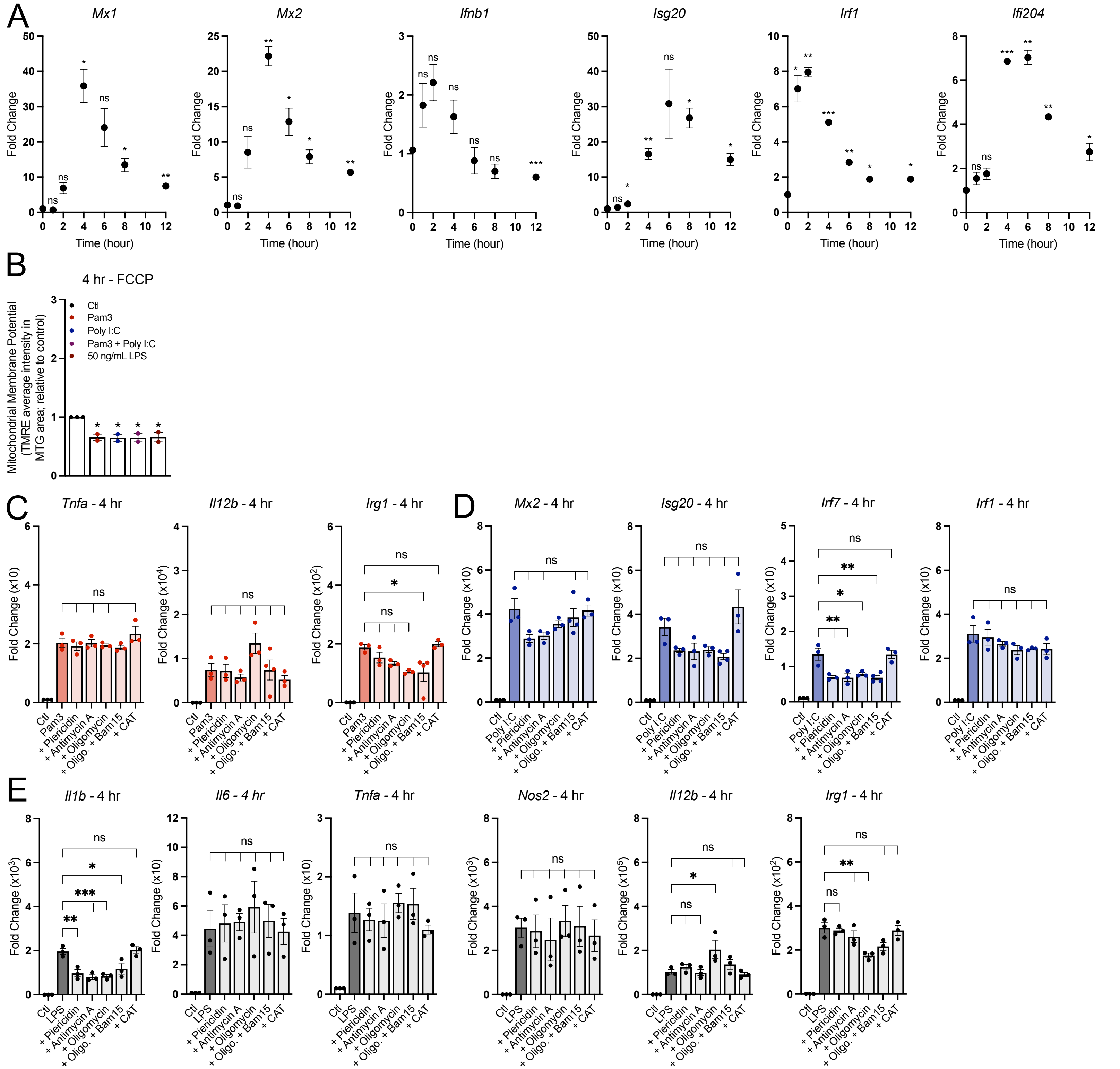


**Supplementary Figure 9: A**) Pro-inflammatory gene expression in BMDMs treated with 50 ng/mL LPS across multiple timepoints relative to control (Ctl) (n = 3). **B**) Bulk membrane potential as measured by TMRE fluorescence per mitochondrial area detected by Mitotracker Green. Data is shown relative to control for BMDMs treated with Pam3, Poly I:C, Pam3 + Poly I:C, or 50 ng/mL LPS for 4 hr. then acutely treated with 2 μM FCCP (n = 4). **C-E**) Pro-inflammatory gene expression of control (Ctl) and BMDMs treated with Pam3 (n = 4-6) (**C**), Poly I:C (**D**), or 10 ng/mL LPS with mitochondrial effector compounds for 4 hr. (n = 3) (**E**). All data are mean ± SEM with statistical analysis conducted on data from biological replicates, each of which included multiple technical replicates, unless otherwise indicated. Statistical analysis for (**A**) was performed as a paired, two-tailed t-test. Statistical analysis for (**B-E**) was performed as an ordinary one-way, ANOVA followed by Tukey's *post hoc* multiple comparisons test.

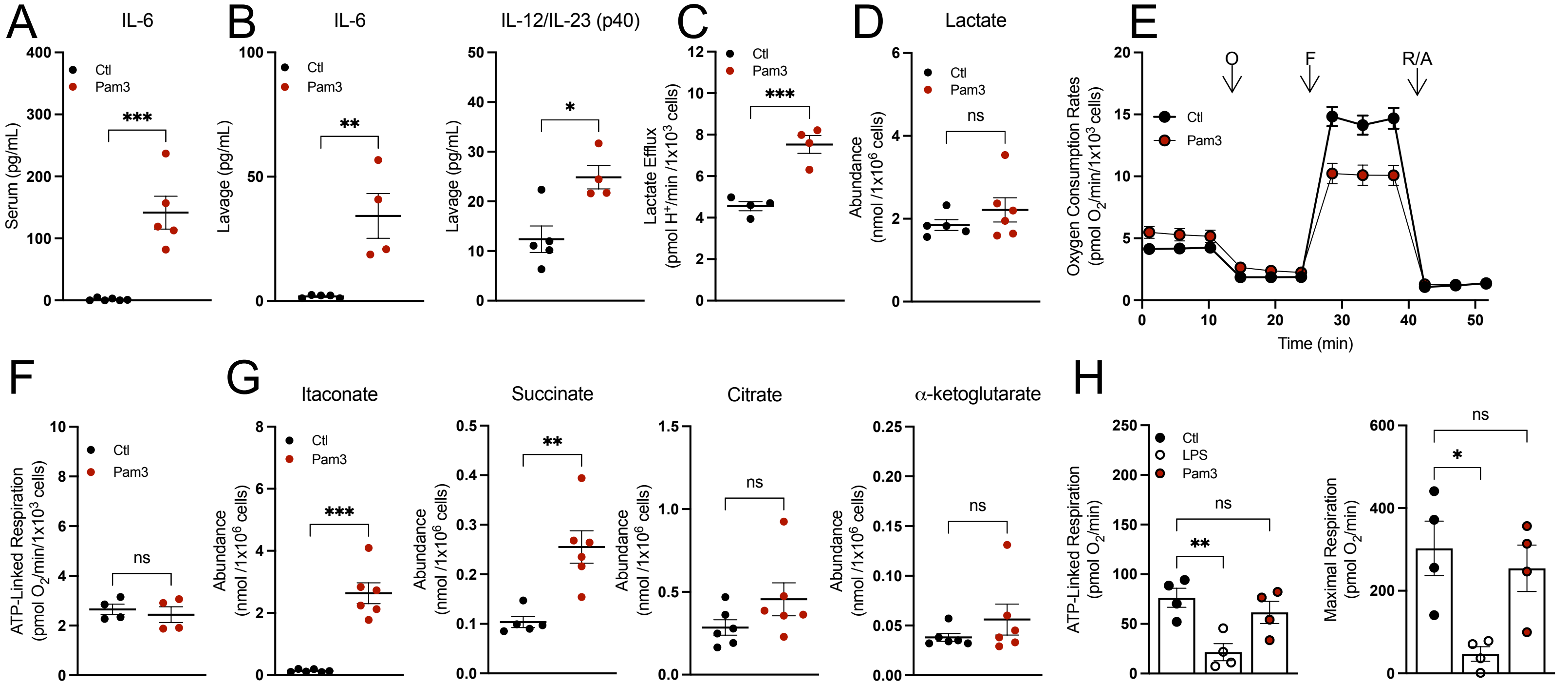


**Supplementary Figure 10: A&B**) Cytokine levels from serum (n = 5-6) (**A**) and lavage fluid from mice intraperitoneally injected with PBS or Pam3 for 24 hr. (n = 4-6) (**B**). **C**) Lactate efflux rates for peritoneal macrophages isolated from mice intraperitoneally injected with PBS or Pam3 for 24 hr. (n = 4). **D**) Intracellular lactate abundance from peritoneal macrophages isolated from mice intraperitoneally injected with PBS or Pam3 for 24 hr. (n = 5-6). **E**) Representative oxygen consumption trace with peritoneal macrophages isolated from mice intraperitoneally injected with PBS or Pam3 for 24 hr. Where not visible, error bars are obscured by the symbol. O, oligomycin; F, carbonyl cyanide-p-trifluoromethoxyphenylhydrazone (FCCP); R/A, rotenone/antimycin A (n = 1 biological replicate with 5 technical replicates). **F**) ATP-Linked respiration for peritoneal macrophages isolated from mice intraperitoneally injected with PBS or Pam3 for 24 hr. (n = 4). **G**) Intracellular itaconate, succinate, citrate and $\alpha$-ketoglutarate abundances from peritoneal macrophages isolated from mice intraperitoneally injected with PBS or Pam3 for 24 hr. (n = 6). **H**) ATP-linked and maximal respiration rates for peritoneal macrophages treated *in vitro* with 50 ng/mL LPS or 50 ng/mL Pam3 for 24 hr. (n = 4). All data are mean ± SEM with statistical analysis conducted on data from biological replicates, each of which included multiple technical replicates, unless otherwise indicated. Statistical analysis for (**A-D**) and (**F-G**) was performed as an an unpaired, two-tailed t-test. Statistical analysis for (**H**) was performed as an ordinary one-way, ANOVA followed by Tukey's *post hoc* multiple comparisons test.
